# Sodium leak channel as a therapeutic target for neuronal sensitization in neuropathic pain

**DOI:** 10.1101/2020.08.17.253534

**Authors:** Donghang Zhang, Wenling Zhao, Jin Liu, Mengchan Ou, Peng Liang, Jia Li, Yali Chen, Daqing Liao, Siqi Bai, Jiefei Shen, Xiangdong Chen, Han Huang, Cheng Zhou

## Abstract

Neuropathic pain affects up to 10% of the total population and no specific target is ideal for therapeutic need. The sodium leak channel (NALCN), a voltage-independent cation channel, mediates the background Na^+^ leak conductance and controls neuronal excitability and rhythmic behaviors. Here, we show that increases of NALCN expression and function in dorsal root ganglion (DRG) and dorsal spinal cord contribute to chronic constriction injury (CCI)-induced neuropathic pain in rodents. NALCN current and neuronal excitability in acutely isolated DRG neurons and spinal cord slices of rats were increased after CCI which were decreased to normal levels by NALCN-siRNA. Accordingly, pain-related symptoms were significantly alleviated by NALCN-siRNA-mediated NALCN knockdown and completely reversed by NALCN-shRNA-mediated NALCN knockdown in rats or by conditional NALCN knockout in mice. Our results indicate that increases in NALCN expression and function contribute to CCI-induced neuronal sensitization; therefore, NALCN may be a novel therapeutic target for neuropathic pain.

## Introduction

Neuropathic pain, caused by injury or disease of nervous system, affects up to 10% of the total population (Colloca et al., 2017). Neuropathic pain encompasses a broad range of clinical symptoms including hyperalgesia, allodynia and spontaneous pain (Gilron et al., 2015). Current therapy for neuropathic pain includes pharmacological, non-pharmacological and interventional treatments, but none of these approaches have a definite target of action (Gilron et al., 2015). Currently, antidepressants (e.g. tricyclic agents and serotonin-norepinephrine reuptake inhibitors) and anticonvulsants (e.g. gabapentin and pregabalin) are the first-line pharmacological treatments for neuropathic pain (Finnerup et al., 2015). However, none of these agents are satisfactory in therapy of neuropathic pain (Colloca et al., 2017).

Several mechanisms have been proposed for the development and maintenance of neuropathic pain, such as aberrant ectopic activity in nociceptors, central neuronal sensitization, impaired descending inhibitory modulation and pathological activation of microglia (Campbell and Meyer, 2006; von Hehn et al., 2012). Although regulation of ion channels contributes to the central neuronal sensitization and hyperactivity in neuropathic pain (Colloca et al., 2017; Kim et al., 2014), the exact causes of neuronal sensitization are not fully clear.

NALCN is a voltage-independent, tetrodotoxin (TTX)-resistant and non-selective cation channel that generates a “leak” inward current under physiological conditions (Cochet-Bissuel et al., 2014; Lu et al., 2007; Ren, 2011). NALCN is widely expressed in the central nervous system and produces a basal Na^+^-leak conductance that regulates neuronal excitability (Lu et al., 2007). NALCN is associated with important nervous system functions such as locomotor behaviors (Xie et al., 2013), respiratory rhythm (Lu et al., 2007) and responsiveness to general anesthetics (Jospin et al., 2007; Singaram et al., 2011; Yang et al., 2020). A recent study indicates that expression of NALCN may contribute to sensory processing by regulating the intrinsic excitability of spinal projection neurons (Ford et al., 2018). The overexpression of NALCN can induce depolarization of resting membrane potential in nociceptors, which leads to pain insensitivity in African highveld mole-rats (Eigenbrod et al., 2019). Additionally, NALCN is also involved in pain processing in thermo-nociceptor of *C. elegans* (Saro et al., 2020). Therefore, NALCN might mediate the neuronal excitability involved in pain sensation and processing. NALCN current can be activated by neuromodulators such as substance P (SP) and neurotensin (Lu et al., 2009; Ren, 2011). SP can activate NALCN *via* the SP receptor (NK1R) in a G protein-independent manner that requires the Src family of kinases and the UNC80 subunit (Ren, 2011). SP is well-known for its important role in pain transmission through NK1R in spinal cord (Todd, 2010), and release of SP is increased in various conditions, including neuropathic pain (Tiwari et al., 2014). Whether NALCN is a target of SP in pain perception is unknown.

In the present study, we aimed to determine whether regulation of NALCN expression and function contributes to neuropathic pain. The results indicate that NALCN expression in DRG and dorsal spinal cord increases with symptom of neuropathic pain. NALCN activity increases with neuronal hyperactivity in DRG and spinal sensory neurons after chronic constriction injury (CCI)-induced neuropathic pain. Knockdown of NALCN expression by NALCN-siRNA in DRG and spinal cord neurons significantly alleviated the mechanical allodynia and thermal hyperalgesia in rats, as well as NALCN current and neuronal excitability. Notably, knockdown of NALCN expression in the DRG and spinal cord neurons by NALCN-shRNA in rats or conditional knockout of NALCN in the DRG and spinal cord neurons in mice completely reversed the development of CCI-induced mechanical allodynia and thermal hyperalgesia. This present study provides novel insights into neuronal sensitization during neuropathic pain by elucidating the involvement of NALCN, which might serve as a therapeutic target.

## Results

### Expression of NALCN mRNA increases after chronic constriction injury (CCI) in rats

The CCI model was established in adult male Sprague-Dawley (SD) rats, and the CCI-ipsilateral hind limb developed mechanical allodynia and thermal hyperalgesia beginning on the first day after CCI and lasting for at least 2 weeks (Fig. 1A). The NALCN mRNA level in the DRG and dorsal spinal cord from the CCI-ipsilateral side was significantly increased compared with the CCI-contralateral side at day 3 (Fig. 1B) and day 14 (Fig. 1C) after CCI. NALCN was widely present in DRG neurons that labeled by neurofilament (NF200, A-fibers), transient receptor potential cation channel subfamily V member 1 (TRPV1, noxious receptor), isolectin B4 (IB4, non-peptidergic neurons) and substance P (SP, peptidergic neurons) (Fig. 1D). Therefore, NALCN was expressed in nearly all neurons of DRG (Fig. 1E). The percentage of NALCN-positive neurons that also expressed SP was greater in DRG of the CCI-ipsilateral side at day 14 after CCI (Fig. 1F). NALCN was also widely expressed in neurons of spinal cord (Fig. 1G). In addition, we also measured age- and tissue-specific expression of NALCN. Similar with adult rats, NALCN mRNA was also abundantly expressed in DRG and dorsal spinal cord in rats at the age of postnatal day 8 (Supplementary fig. 1A). Compared with the spinal cord, the level of NALCN mRNA was much lower in the heart of both neonatal and/or adult rats (Supplementary fig. 1B).

**Fig. 1.**
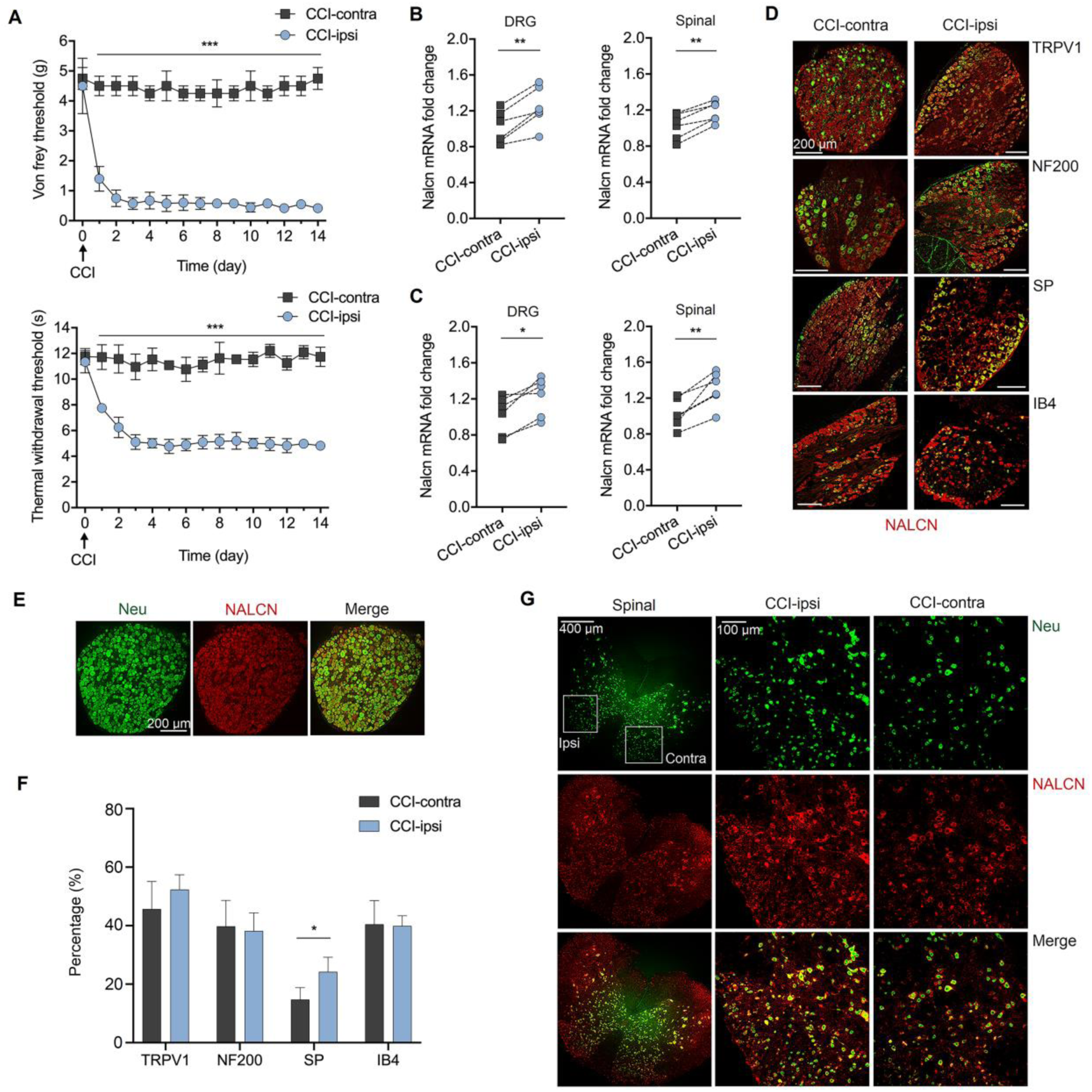
Expression of NALCN increases after CCI. (**A**) Mechanical allodynia and thermal hyperalgesia were developed in CCI-ipsilateral paw compared to CCI-contralateral side since the second day after CCI until at least day 14 (n=8). (**B** and **C**) NALCN mRNA in DRG and dorsal spinal cord from CCI-ipsilateral side were significantly increased compared to CCI-contralateral side at both day 3 (B, n = 6) and day 14 (C, n = 6) after CCI. (**D**) DRG neurons from the CCI-contralateral and CCI-ipsilateral side were double labeled with NALCN (red) and SP, NF200, TRPV1 and IB4 (green) respectively. (**E**) DRG neurons were double labeled with NALCN and NeuN, indicating NALCN was present in almost all the neurons. (**F**) Percentage of SP-positive neurons that express NALCN was significantly increased in the CCI-ipsilateral side compared to the CCI-contralateral side at day 14 (n = 5-7 sections from 3 rats). (**G**) Neurons in spinal cord were double labeled with NALCN and NeuN, indicating NALCN was widely expressed and higher fluorescent intensity was found in the CCI-ipsilateral side compared to the CCI-contralateral side at day 14. The middle panel (CCI-ipsi) and right panel (CCI-contra) are enlarged from the white square of the left panel. Data are present as mean ± SEM. A by two-way ANOVA; B, C by paired two-tailed student’s t-test; F by unpaired two-tailed student’s t-test. * *P* < 0.05, ** *P* < 0.01 and *** *P* < 0.001.

### Increased expression of NALCN contributes to the hyperactivity of sensory neurons after CCI in rats

Neuronal sensitization and hyperactivity is characterized by exaggerated responses to the normal stimuli (Campbell and Meyer, 2006). Here we determined whether increased expression of NALCN contributes to the hyperactivity of sensory neurons after CCI. NALCN-siRNA or control-siRNA was intrathecally and intraneurally injected immediately after CCI in the rats at age of postnatal day 5 (P5) (Fig. 2A). Whole-cell patch-clamp recordings were performed on acutely isolated DRG neurons and spinal cord slices 2-3 days later. For the rats received control-siRNA, dorsal spinal cord and DRG neurons from the CCI-ipsilateral side were hyperactive (Fig. 2C-E, G-I left panel), evidenced by depolarized resting membrane potential (RMP) (Fig. 2C left panel for spinal cord and Fig. 2G left panel for DRG), decreased rheobase (Fig. 2D left panel for spinal cord and Fig. 2H left panel for DRG), and decreased input resistance (R_input_) (Fig. 2E left panel for spinal cord and Fig. 2I left panel for DRG), compared to the CCI-contralateral side. With NALCN-siRNA treatment, the profiles of neuronal activities were similar in the neurons between the CCI-ipsilateral side and CCI-contralateral side (Fig. 2C-E, G-I right panel). These results indicate that neurons are hyperactive in the DRG and dorsal spinal cord of CCI-ipsilateral side, at least partly contributed by increased expression of NALCN. As similar with the results in adult rats, RT-PCR results showed that the NALCN mRNA level in the DRG and dorsal spinal cord from the CCI-ipsilateral side was significantly increased compared with the CCI-contralateral side at day 3 after CCI in P5 rats (Fig. 2J). As a control experiment, NALCN mRNA level was significantly decreased in the DRG by NALCN-siRNA (Fig. 2K, left) but not control-siRNA (Fig. 2K, right), and NALCN mRNA level was significantly decreased in the dorsal spinal cord by NALCN-siRNA when compared to control-siRNA (Fig. 2L) at 3 days after injection. Furthermore, NALCN mRNA was decreased in the DRG (Fig. 2M, left panel) and not increased in dorsal spinal cord (Fig. 2M, right panel) on the CCI-ipsilateral side compared with the CCI-contralateral side at 3 days after CCI in rat pups that received NALCN-siRNA. The rats that underwent CCI at P5 exhibited mechanical allodynia (Fig. 2N) and thermal hyperalgesia (Fig. 2O) when tested at P30.

**Fig. 2.**
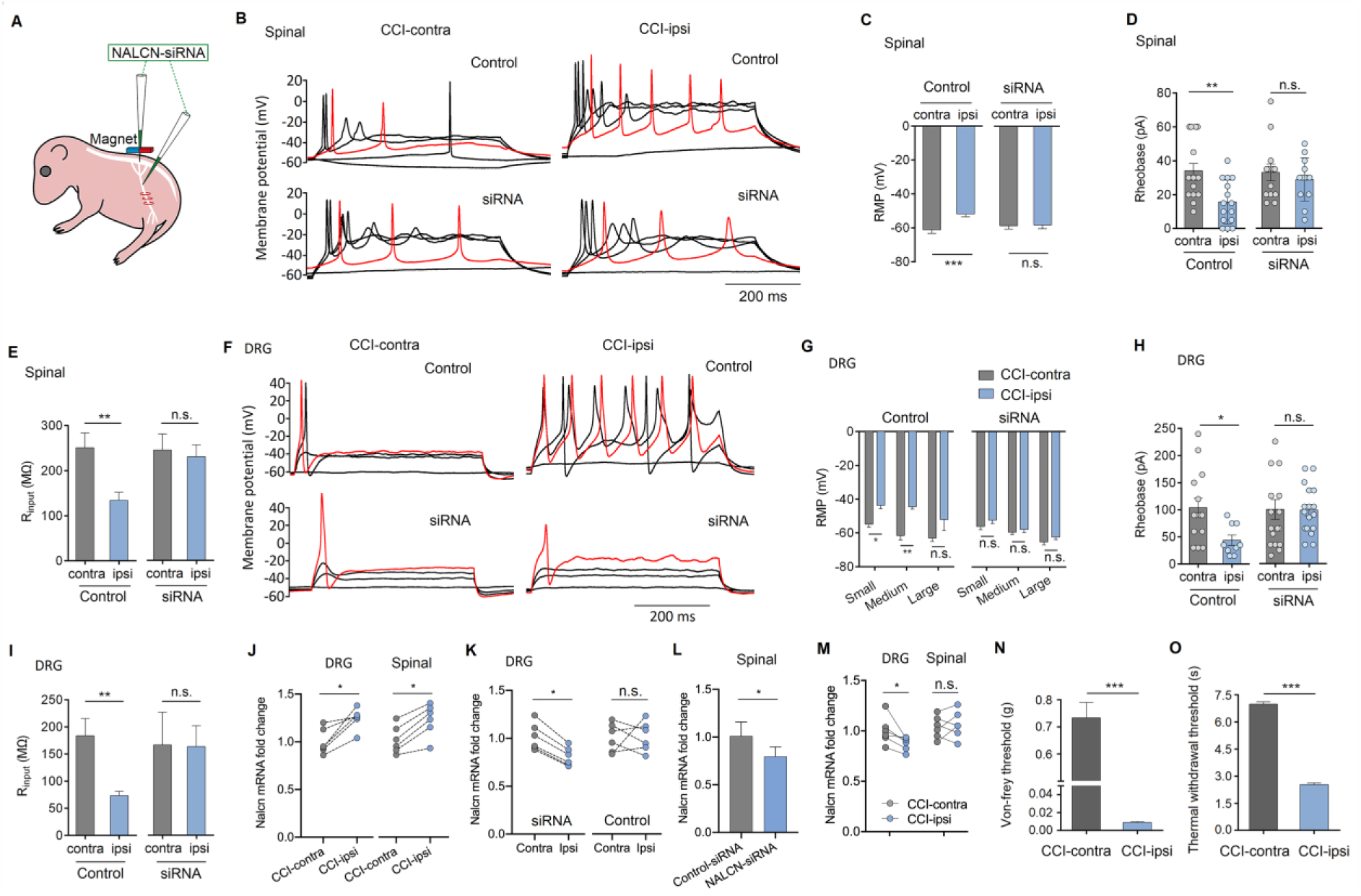
NALCN enhances the excitability of sensory neurons after CCI in rats. (**A**) Representative cartoon of intraneural and intrathecal injection of NALCN-siRNA. (**B**) Representative traces of action potentials in the neurons of dorsal spinal cord. Red trace is the action potentials when receiving the same current injection. (**C** to **E**) Excitability of spinal neurons was compared between CCI-contralateral and CCI-ipsilateral side based on rest membrane potential (RMP) (C, left panel, n = 20-29; C, right panel, n=13), rheobase (D, left panel, n = 15-16; D, right panel, n = 13), and input resistance (R_input_) (E, left panel, n = 11; E, right panel, n = 12-13). (**F**) Representative traces of action potentials in the DRG neurons. Red trace is the action potentials when receiving the same current injection. (**G** to **I**) Excitability of DRG neurons was compared between CCI-contralateral and CCI-ipsilateral side based on rest membrane potential (RMP) (G, left panel, n = 4-17; G, right panel, n = 10-15), rheobase (H, left, n=9-14; H, right, n=17), and input resistance (R_input_) (I, left panel, n=9-11; I, right panel, n = 12). (**J**) NALCN mRNA level in the DRG (left panel, n=6) and dorsal spinal cord (right panel, n=6) from the CCI-ipsilateral side was significantly increased compared with the CCI-contralateral side at day 3 after CCI model in P5 rats. (**K**) NALCN mRNA level was decreased in the DRG (K, left panel, n=6) by NALCN-siRNA but not control-siRNA (K, right panel, n=6) at day 3 after injection. (**L**) NALCN mRNA level was decreased in the dorsal spinal cord (n=6) by NALCN-siRNA when compared to control-siRNA at day 3 after injection. (**M**) NALCN mRNA was decreased in the DRG (left panel, n=6) and not increased in dorsal spinal cord (right panel, n=6) by NALCN-siRNA on the CCI-ipsilateral side compared with the CCI-contralateral side at day 3 after CCI in P5 rats. (**N** and **O**) Mechanical allodynia (N, n = 12) and thermal hyperalgesia (O, n = 12) were observed at P30 in the CCI-ipsilateral paws of the rats received CCI at the age of P5. Data are present as mean ± SEM. C, D, E, G, H, I right panel, L by unpaired two-tailed student’s t-test; I left panel by two-tailed Mann-Whitney test; J, K, M, N, O by paired two-tailed student’s t-test. * *P* < 0.05, ** *P* < 0.01 and *** *P* < 0.001. n.s.: no significance.

### NALCN current is functionally increased after CCI in rats

The NALCN-mediated holding current (*I*_holding_) was recorded at a holding potential of −60 mV, and measured by replacing normal extracellular Na^+^ with NMDG (Fig. 3A, G, M, P). For the rats received control-siRNA, the *I*_holding_ was significantly increased in the neurons from dorsal spinal cord (Fig. 3C) and DRG (Fig. 3O) of CCI-ipsilateral side, compared to CCI-contralateral side. NALCN-siRNA treatment diminished the functional difference between the CCI-ipsilateral and CCI-contralateral sides in both dorsal spinal cord (Fig. 3I) and DRG (Fig. 3R) neurons.

**Fig. 3.**
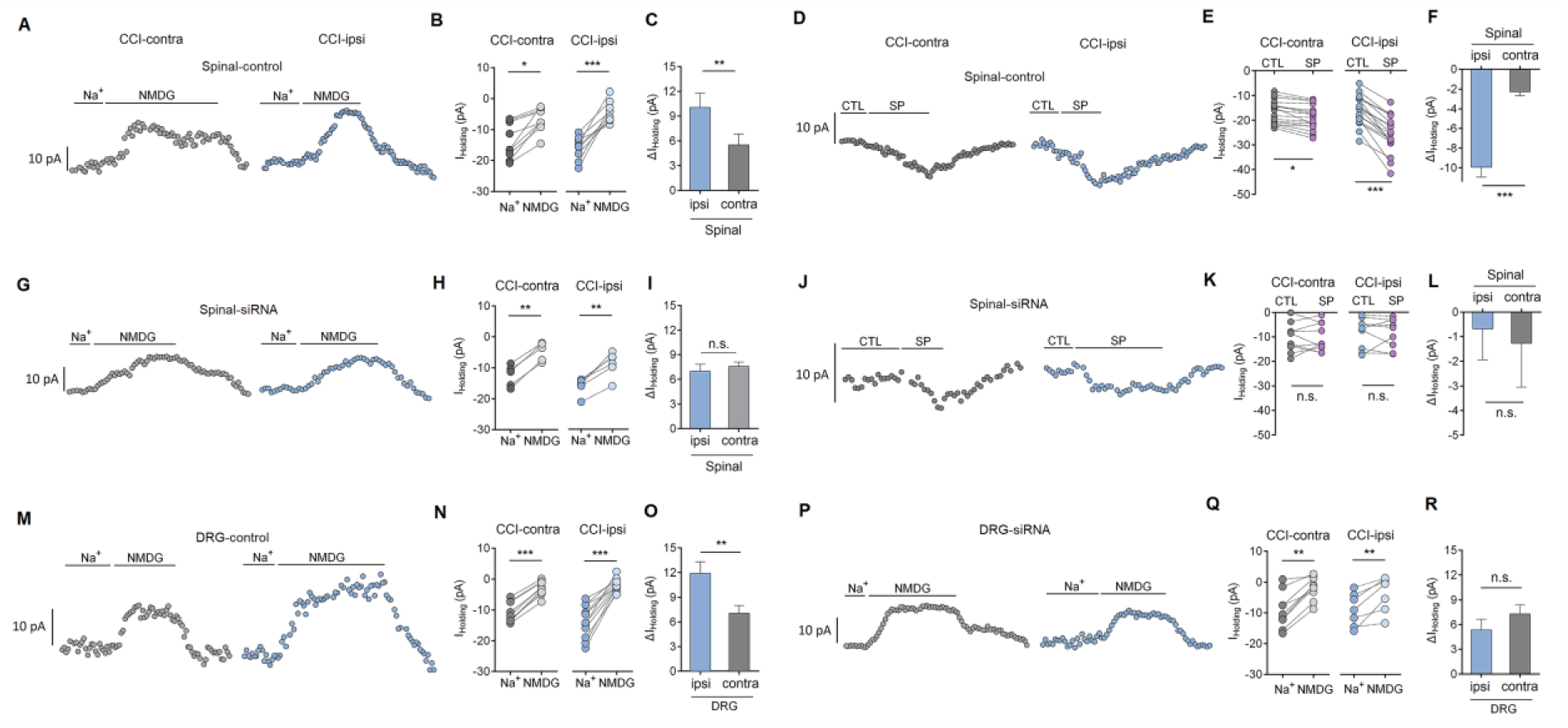
NALCN current and responses increase after CCI in rats. (**A, G, M, P**) The representative real-time analysis of holding currents in neurons between CCI-contralateral and CCI-ipsilateral side. NALCN current was recorded at holding potential of −60 mV and extracted by replacing extracellular Na^+^ with NMDG. (**B, H, N, Q)** Holding currents were recorded before and after replacement with NMDG. (**C, I, O, R)** Changed holding currents were compared between CCI-contralateral and CCI-ipsilateral side after replacement with NMDG (C, n = 9; I, n = 5-6; O, n = 9-11; R, n=7-9). (**D** and **J**) The representative real-time analysis of holding currents in neurons between CCI-contralateral and CCI-ipsilateral side. SP evoked holding currents were recorded at holding potential of −60 mV. (**E** and **K**) Holding currents were recorded before and after perfusion of SP. (**F** and **L**) Right panel is changed holding currents after perfusion of SP (F, n = 17; L, n = 9-10). Data are present as mean ± SEM. C, F, I, O, R by unpaired two-tailed student’s t-test; L by two-tailed Mann-Whitney test. ** *P* < 0.01 and *** *P* < 0.001. n.s.: no significance.

The neuropeptide substance P (SP) can enhance NALCN conductance *via* the neurokinin-1 receptor (NK1R) (Lu et al., 2009). The SP-evoked *I*_holding_ was also recorded and significantly larger in dorsal spinal cord (Fig. 3F) neurons of the CCI-ipsilateral side than in the neurons of CCI-contralateral side from the rats received control-siRNA. For the rats received NALCN-siRNA, no difference was found in neurons between the CCI-ipsilateral and CCI-contralateral sides from dorsal spinal cord (Fig. 3L). These results indicate that the NALCN-mediated *I*_holding_ is increased after CCI, which can strengthen neuronal responses to SP.

Gd^3+^ is a non-selective blocker of NALCN. Gd^3+^-mediated inhibition of the *I*_holding_ was significantly greater in DRG (Fig. 4B) and dorsal spinal cord (Fig. 4E) neurons on the CCI-ipsilateral side than CCI-contralateral side in the rats received control-siRNA. For the rats received NALCN-siRNA, no difference was found between the CCI-ipsilateral and CCI-contralateral side in DRG (Fig. 4H) and dorsal spinal cord (Fig. 4K) neurons. Current-voltage (I-V) curves of leak currents (−100 to −40 mV) were compared between the CCI-ipsilateral and CCI-contralateral sides (Fig. 4C, F, I, L, left panel).

**Fig. 4.**
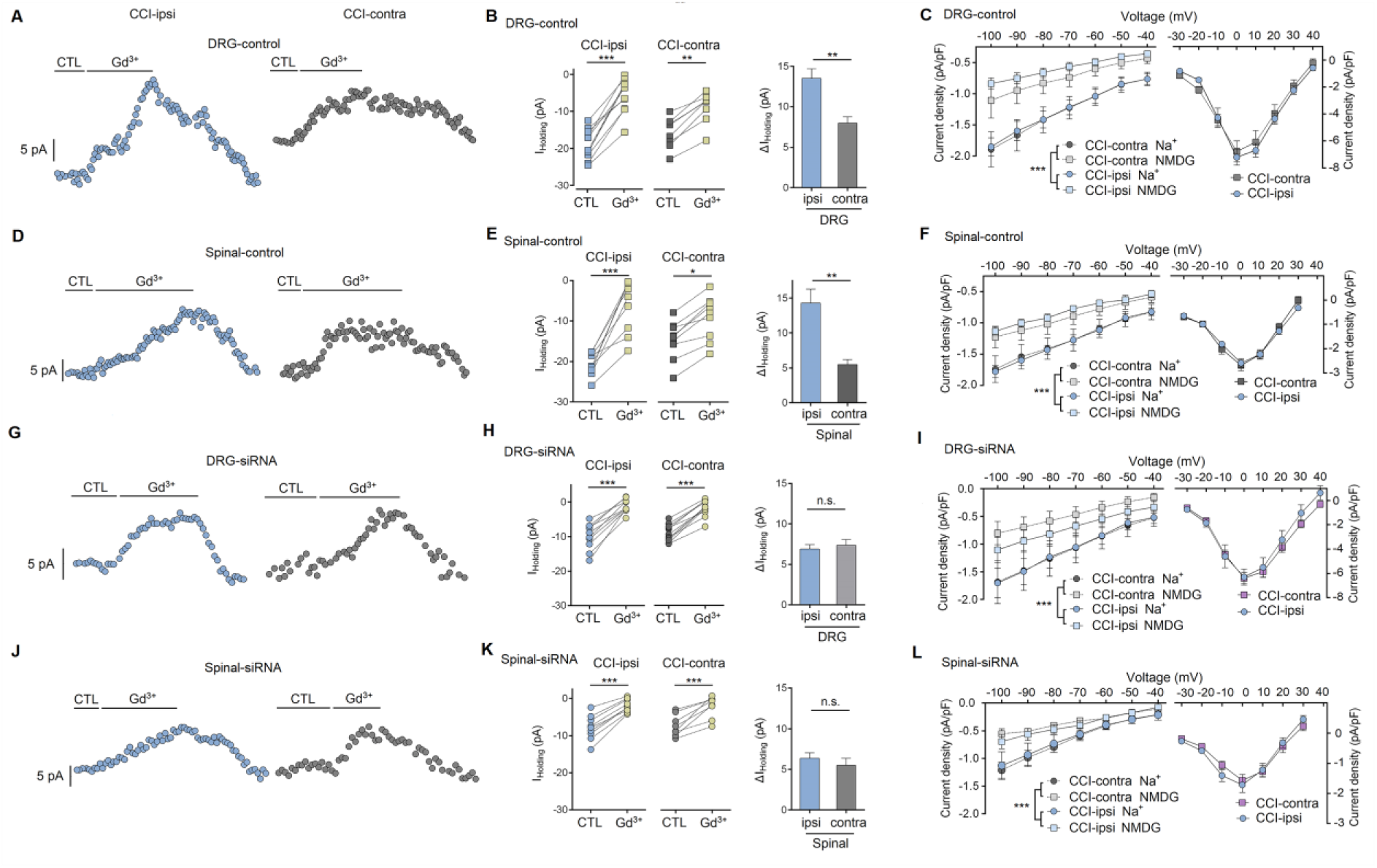
Gd^3+^-inhibited holding currents and I-V (Current-voltage) currents were recorded after CCI in rats. (**A, D, G, J**) Representative real-time analysis of Gd^3+^-inhibited holding currents. Sensory neurons of DRG and dorsal spinal cord (Lamina I-II) were recorded at holding potential of −60 mV. (**B, E, H, K**) Left panel is the holding currents before and after perfusion of Gd^3+^ between. Right panel is the changed holding currents after perfusion of Gd^3+^ (B, n = 9-11; H, n = 13-14; E, n = 9; K, n = 9-10). (**C, F, I, L**) The neurons of acute isolated DRG and dorsal spinal cord were recorded and compared between the rats received control-siRNA (C and F) and NALCN-siRNA (I and L). I-V curve from voltages of −100 mV to −40 mV was compared after replacement of extracellular Na^+^ with NMDG between CCI-contralateral and CCI-ipsilateral side (C, left panel, n = 9-12; I, left panel, n=10-11; F, left panel, n = 11-12; L, left panel, n = 5-6). I-V curves since voltage of −30 mV was compared between CCI-contralateral and CCI-ipsilateral side. TTX-resistant Na_v_ was activated within these voltage levels. The density of TTX-R *I*_Na_ was unchanged at 2-3 days after CCI between CCI-ipsilateral side and CCI-contralateral side (C, right panel, n = 8-15; I, right panel, n = 10-11; F, right panel, n = 11-12; L, right panel, n = 6-7). Data are present as mean ± SEM. B, E, H and K by unpaired two-tailed student’s t-test; C, F, I and L by two-way ANOVA. * *P* < 0.05, ** *P* < 0.01 and *** *P* < 0.001. n.s.: no significance.

Na^+^-mediated currents were measured by replacing normal extracellular Na^+^ with NMDG. Na^+^-mediated leak current was increased in the neurons from the CCI-ipsilateral side in the rats received control-siRNA, compared with CCI-contralateral side (Fig. 4C, F, left panel). NALCN-siRNA treatment decreased the Na^+^-mediated leak current in the neurons from the CCI-ipsilateral side than CCI-contralateral side (Fig. 4I, L, left panel).

Regulation of voltage-gated sodium channels (Na_v_) is also important for neuronal excitability (Colloca et al., 2017; Tibbs et al., 2016). Tetrodotoxin (TTX) was perfused when recoding NALCN-mediated leak currents. Therefore, we also analysed TTX-resistant (TTX-R) Na_v_ currents (*I*_Na_). The density of TTX-resistant (TTX-R) Na_v_ currents (*I*_Na_) was unchanged (Fig. 4C, F, right panel). NALCN-siRNA did not affect the density of TTX-R *I*_Na_ (Fig. 4I, L, right panel). These results indicate that TTX-R Na_v_ may not be involved in the initiation phase of rapid neuronal sensitization after CCI, and the NALCN-siRNA here is selective between NALCN and TTX-R Na_v_. However, the contribution of Na_v_ cannot be excluded for maintenance of neuronal hyperactivity later (Colloca et al., 2017; Tibbs et al., 2016).

### Knockdown expression of NALCN in the DRG and spinal cord reduces SP-induced pain behaviors in rats

Intrathecal injection of SP can induce acute pain-related behaviors, including biting and scratching of the bilateral skin (Hylden and Wilcox, 1981). To test whether NALCN was involved in SP-induced aversive response, NALCN-siRNA or control-siRNA was intrathecally and intraneurally injected into the subarachnoid space and sciatic nerve in adult rats (Fig. 5A). Normal sensory function including mechanical and thermal sensation was not affected compared to baseline when measured at day 3 after NALCN-siRNA injection (Fig. 5B). Interestingly, knockdown the expression of NALCN in the DRG and spinal cord by NALCN-siRNA significantly reduced the total time that the rats spent biting and scratching within the first 5 min after intrathecal injection of SP (Fig. 5C). Decrease of NALCN mRNA was confirmed by RT-PCR after the behavioral tests (3 days after injection of NALCN-siRNA) (Fig. 5D).

**Fig. 5.**
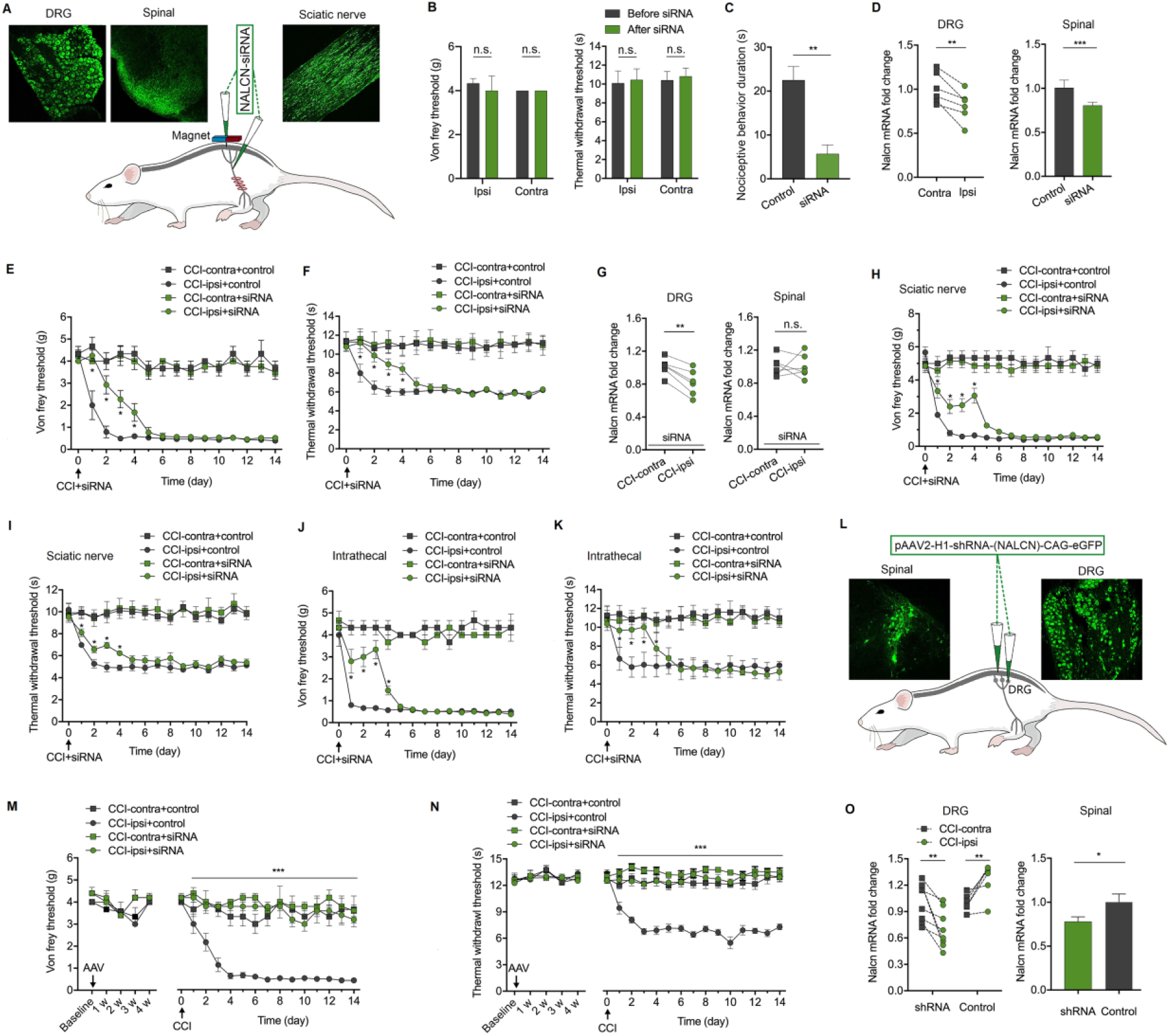
Knockdown the expression of NALCN in DRG and spinal cord alleviates pain behaviors induced by CCI in rats. (**A**) Representative cartoons of NALCN-siRNA injection (bottom). Alexa Fluor 488-positive fluorescence was present in DRG, dorsal spinal cord and sciatic nerve after intraneural and intrathecal injection of NALCN-siRNA. (**B**) Normal sensory function was unchanged by NALCN-siRNA in von Frey (left panel, n = 6) or thermal test (right panel, n = 6), compared to baseline. (**C**) Intrathecal injection of SP induced pain related behaviors in rats including biting and scratching at the abdomen and hind portions of the body. The duration of biting and scratching of rats within the first 5 min after injection of SP was diminished in the rats received NALCN-siRNA (n = 6). (**D**) The level of NALCN mRNA was significantly decreased in DRG at day 3 after injection of NALCN-siRNA (left panel, n = 6) but not control-siRNA. The level of NALCN mRNA was significantly decreased in dorsal spinal cord (right panel, n = 6) at day 3 after injection of NALCN-siRNA as compared to control-siRNA. (**E** and **F**) NALCN-siRNA significantly alleviated mechanical allodynia (E, n = 6 for control-siRNA group, n = 8 for NALCN-siRNA group, \**P* < 0.05 vs. control-siRNA) and thermal hyperalgesia (F, n = 6 for control-siRNA group, n = 8 for NALCN-siRNA group, \**P* < 0.05 vs. control-siRNA) at day 1-4 after CCI. (**G**) NALCN mRNA was decreased in DRG (left panel, n = 6) and not increased in dorsal spinal cord (right panel, n = 6) at day 3 after CCI by NALCN-siRNA. (**H** and **I**) Intraneural (sciatic nerve) injection of NALCN-siRNA alleviated the CCI-induced mechanical allodynia (H, n = 7, \**P* < 0.05 vs. control-siRNA) and thermal hyperalgesia (I, n=7, \**P* < 0.05 vs. control-siRNA) after CCI. (**J** and **K**) Intrathecal injection of NALCN-siRNA significantly alleviated the CCI-induced mechanical allodynia (J, n = 7, \**P* < 0.05 vs. control-siRNA) and thermal hyperalgesia (K, n = 7, \**P* < 0.05 vs. control-siRNA) after CCI. (**L**) Representative cartoons of pAAV2-H1-shRNA-(NALCN)-CAG-eGFP injection (bottom). GFP-positive fluorescence was found in DRG and dorsal spinal cord 4 weeks after DRG injection and intrathecal injection of pAAV2-H1-shRNA-(NALCN)-CAG-eGFP (AAV-NALCN-shRNA) or pAAV2-scrambled-CAG-eGFP virus (AAV-scrambled-shRNA) (top). (**M** and **N**) Normal sensory function in von Frey (M, left panel, n = 6 for AAV-scrambled-shRNA group, n = 10 for AAV-NALCN-shRNA group) or thermal test (N, left panel, n = 6 for AAV-scrambled-shRNA group, n=10 for AAV-NALCN-shRNA group) was unchanged during the 4 weeks after AAV-NALCN-shRNA injection, compared to AAV-scrambled-shRNA or baseline. NALCN-shRNA alleviated mechanical allodynia (M, right panel, n = 6 for AAV-scrambled-shRNA group, n=10 for AAV-NALCN-shRNA group) and thermal hyperalgesia (N, right panel, n = 6 for AAV-scrambled-shRNA group, n = 10 for AAV-NALCN-shRNA group) throughout after CCI, compared to scrambled shRNA. (**O**) NALCN mRNA was decreased in DRG in the CCI-ipsilateral side of AAV-NALCN-shRNA group but increased in DRG in the CCI-ipsilateral side of AAV-scrambled-shRNA group at day 14 after CCI (left panel, n = 6 for AAV-scrambled-shRNA group, n = 8 for AAV-NALCN-shRNA group). NALCN mRNA was decreased in dorsal spinal cord of AAV-NALCN-shRNA group as compared to AAV-scrambled-shRNA group at day 14 after CCI (right panel, n = 6 for AAV-scrambled-shRNA group, n = 8 for AAV-NALCN-shRNA group). Data are present as mean ± SEM. B left panel, C, G and O left panel by paired two-tailed student’s t-test; B right panel and O right panel by unpaired two-tailed student’s t-test; D by two-tailed Mann-Whitney test; E, F, H, I, J, K, M and N by two-way ANOVA; * *P* < 0.05, ** *P* < 0.01 and *** *P* < 0.001. n.s.: no significance.

### Knockdown expression of NALCN in the DRG and spinal cord alleviates the mechanical allodynia and thermal hyperalgesia after CCI in rats

To test the therapeutic role of NALCN in neuropathic pain, NALCN-siRNA or control-siRNA was intrathecally and intraneurally injected immediately after CCI. Mechanical allodynia (Fig. 5E) and thermal hyperalgesia (Fig. 5F) of the hind limb in CCI-ipsilateral side were significantly alleviated by NALCN-siRNA from day 1 to day 4 after CCI. The NALCN mRNA was decreased in the DRG (Fig. 5G, left panel) and not increased in dorsal spinal cord (Fig. 5G, right panel) by NALCN-siRNA on the CCI-ipsilateral side compared with the CCI-contralateral side at day 3 after CCI. The region-specific role of NALCN in neuropathic pain was also investigated. Intraneural (Fig. 5H-I) or intrathecal (Fig. 5J-K) injection of NALCN-siRNA also relieved CCI-induced hyperalgesia, indicating that preventing up-regulation of NALCN in DRG and/or spinal cord can suppress the initiation of neuropathic pain *in vivo*.

NALCN-siRNA is degraded rapidly and only have a short duration of action after injection. To test the role of NALCN in maintenance of neuropathic pain, pAAV2-H1-shRNA-(NALCN)-CAG-eGFP (AAV-NALCN-shRNA) or pAAV2-scrambled-CAG-eGFP (AAV-scrambled-shRNA) virus was injected into DRG and subarachnoid space of adult rats (Fig. 5L). The knockdown efficiency of AAV-NALCN-shRNA was confirmed by RT-PCR at 4 weeks after injection (Supplementary fig. 2A). CCI-induced mechanical allodynia (Fig. 5M) and thermal hyperalgesia (Fig. 5N) were completely reversed to normal throughout by the AAV-NALCN-shRNA but not the AAV-scrambled-shRNA. Accordingly, NALCN mRNA level was also significantly decreased both in the DRG (Fig. 5O, left) and the dorsal spinal cord (Fig. 5O, right) by AAV-NALCN-shRNA at day 14 after CCI.

### The cAMP-PKA pathway regulates NALCN expression in CCI-induced neuropathic pain in rats

To determine whether the cAMP-PKA pathway regulates expression of NALCN in CCI-induced neuropathic pain, a cAMP (SQ22536) or PKA inhibitor (H89) was intrathecally injected for 3 consecutive days after CCI. CCI-induced mechanical allodynia (Supplementary fig. 3A) and thermal hyperalgesia (Supplementary fig. 3B) were significantly improved by SQ22536 from day 1 to day 4 or by H89 from day 1 to day 3 after CCI. The NALCN mRNA level was unchanged in the CCI-ipsilateral side compared to the CCI-contralateral side at day 3 after CCI (Supplementary fig. 3C-D). These results indicate that the cAMP-PKA pathway may regulate the expression of NALCN in CCI-induced neuropathic pain.

### NALCN is abundantly and widely expressed in mice DRG neurons by single-cell RNA sequencing

The re-analysis was based on the data resource of single-cell RNA sequencing (L4-L6 DRG) in adult mice (Usoskin et al.). Here we subdivided cells into 5 categories (Fig. 6A), including peptidergic nociceptors (PEP), non-peptidergic nociceptors (NP), neurofilament containing (NF), tyrosine hydroxylase containing (TH) and non-neuronal clusters with distinct expressional profiles (Supplementary fig. 4) according to the well-known cell-type-specific marker genes: *Nefh*, *Pvalb*, *Tac1*, *Ntrk1*, *Calca*, *Mrgprd*, *P2rx3*, *Th, B2m, Vim, Col6a2* (Usoskin et al.). The cell numbers and the percentage of each category in this analysis was showed in Fig. 6A (right panel). We compared the expression levels of NALCN mRNA with well-known ion channels that expressed in DRG nociceptors, including TRPV1 and the major voltage-gated sodium channels (Na_v_) (Fig. 6B). The results demonstrated that NALCN mRNA was commonly expressed in all neuronal clusters of DRG (Fig. 6B). Compared to NALCN which commonly expressed in all neuronal clusters, TRPV1 was mainly expressed in NP (non-peptidergic nociceptors) and PEP (peptidergic nociceptors) clusters with similar amount to NALCN (Fig. 6C).

**Fig. 6.**
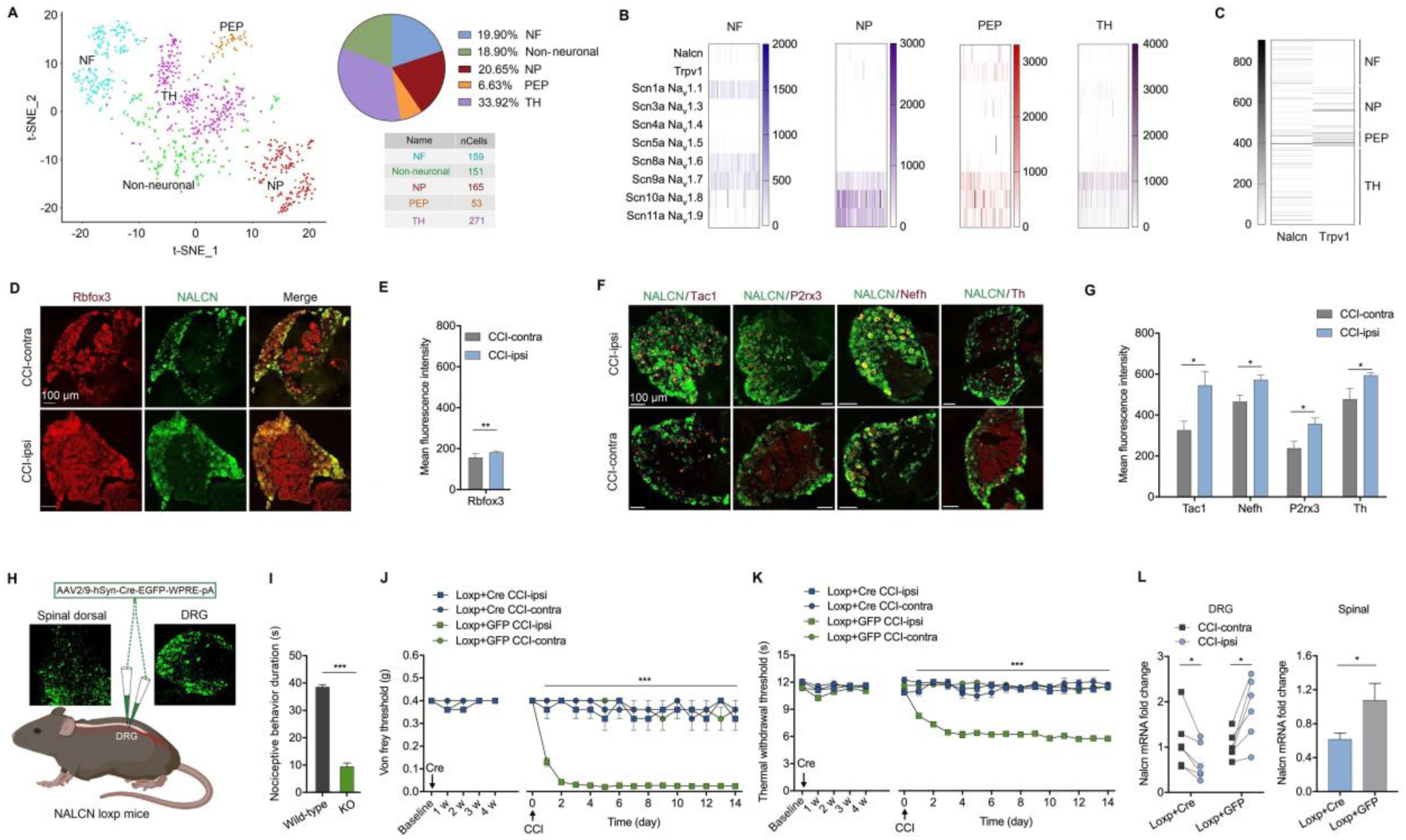
NALCN is abundantly expressed in mice DRG neurons by single-cell RNA sequencing and conditional NALCN knockout in mice DRG and spinal cord neurons completely prevented CCI-induced pain behaviors. (**A**) t-Distributed stochastic neighbor embedding showed 5 major cell populations including PEP, NP, NF, TH and Non-neuronal clusters *via* projection of 799 single-cell transcriptomes from mouse lumbar DRGs. (**B**) Heatmap showed the expression levels of NALCN, TRPV1 and major voltage-gated sodium channels in NF, TH, PEP, NP clusters of DRG, respectively. Numbers reflect the number of UMI detected for the specified gene for each cell. (**C**) Heatmap showed the comparison of expression levels between NALCN and TRPV1 in all cells of DRG. Numbers reflect the number of UMI detected for the specified gene for each cell. (**D** and **E**) Representative images (**C**) and quantifications (E, n = 6-8 sections from three mice) showed the mRNA expression of NALCN in DRG neurons using RNAscope. (**F** and **G**) Representative images (F) and quantifications (G, n = 4-6 sections from three mice) showed the mRNA expression of NALCN in four different subtypes of DRG using RNAscope. (**H**) Representative cartoons of AAV2/9-hSyn-Cre-EGFP-WPRE-pA injection (bottom). GFP-positive fluorescence was found in DRG and dorsal spinal cord 4 weeks after DRG and intrathecal injection of AAV2/9-hSyn-Cre-EGFP-WPRE-pA (AAV-Cre) or AAV2/9-hSyn-EGFP-WPRE-pA (AAV-GFP) (top). (**I**) Knockout of NALCN in the DRG and spinal cord in NALCN^f/f^ mice significantly reduced the total time of biting and scratching within the first 5 min after injection of SP (n=3). (**J** and **K**) Normal sensory function in von Frey (J, left panel, n = 6) or thermal test (K, left panel, n = 6) was unchanged in NALCN^f/f^ mice during 4 weeks after AAV-Cre or AAV-GFP injection as compared or baseline. (**J** and **K**) NALCN knockout completely inhibited the development of mechanical allodynia (J, right panel, n = 6) and thermal hyperalgesia (K, n = 6, right panel) throughout after CCI. (**L**) NALCN mRNA was decreased in DRG from the CCI-ipsilateral side of NALCN^f/f^ mice (L, left panel, n = 6) that received AAV-Cre, but increased in DRG from the CCI-ipsilateral side of NALCN^f/f^ mice (L, left panel, n = 6) that received AAV-GFP as compared to the CCI-ipsilateral side at day 14 after CCI. NALCN mRNA was decreased in the spinal cord of NALCN^f/f^ mice that received AAV-Cre as compared to NALCN^f/f^ mice that received AAV-GFP at day 14 after CCI (L, right panel, n = 6). Data are present as mean ± SEM. E and G by unpaired two-tailed student’s t-test; J and K by two-way ANOVA; L left panel by paired two-tailed student’s t-test; L right panel by two-tailed Mann-Whitney test. * *P* < 0.05, ** *P* < 0.01 and *** *P* < 0.001. n.s.: no significance.

### Expression of NALCN mRNA increases in all neuronal clusters of DRG after CCI in mice

RNAscope technique was used to map the mRNA expression of NALCN in DRG after CCI in mice. The expression of NALCN mRNA increased in DRG neurons with *Rbfox3* (neuron marker) probe at day 14 after CCI (Fig. 6D-E). Then we measured the expression of NALCN mRNA in four neuronal clusters of DRG, including PEP, NP, NF and TH with *Tac1*, *P2rx3*, *Nefh* and *Th* probe, respectively (Usoskin et al.). The results indicated that the expression of NALCN mRNA increased in all the four neuronal clusters of DRG in the CCI-ipsilateral side compared to CCI-contralateral side at 14 days after CCI (Fig. 6F-G).

### Conditional knockout of NALCN in the DRG and spinal cord completely prevented the development of CCI-induced mechanical allodynia and thermal hyperalgesia in mice

To further demonstrate that NALCN was a critical therapeutic target for neuropathic pain, we employed a genetic knockout strategy using the Cre-loxP-mediated recombination system to conditionally delete NALCN from DRG and spinal cord neurons: AAV2/9-hSyn-Cre-EGFP-WPRE-pA (AAV-Cre) or AAV2/9-hSyn-EGFP-WPRE-pA (AAV-GFP) was injected into DRG (L4-L5) and intrathecally injected into subarachnoid space in the NALCN^f/f^ (NALCN-loxP) mice or NALCN^+/+^ mice (Fig. 6H). The knockout efficiency of AAV-Cre was confirmed by RT-PCR (Supplementary fig. 2B) and RNAscope (Supplementary fig. 5A) 4 weeks later. Consistent with the results in rats, knockout of NALCN in the DRG and spinal cord in NALCN^f/f^ mice significantly reduced the total time that spent biting and scratching within the first 5 min after intrathecal injection of SP (Fig. 6I). Normal sensory function including mechanical (Fig. 6J, left) and thermal (Fig. 6K, left) sensation was not affected in NALCN^f/f^ mice after injection of AAV-Cre until four weeks later. CCI was performed 4 weeks after injection of AAV-Cre and the results showed that CCI-induced mechanical allodynia (Fig. 6J, right) and thermal hyperalgesia (Fig. 6K, right) were completely prevented in the NALCN^f/f^ mice that received AAV-Cre. Accordingly, NALCN mRNA level was significantly decreased in both the DRG (Fig. 6L, left) and the dorsal spinal cord (Fig. 6L, right) of NALCN^f/f^ + AAV-Cre mice at day 14 after CCI. Injection of AAV-Cre in NALCN^+/+^ mice did not produce any affection on the CCI-induced mechanical allodynia (Supplementary Fig. 6A) and thermal hyperalgesia (Supplementary Fig. 6B).

## Discussion

Effective treatment for neuropathic pain is currently unavailable due to the ambiguity of its exact etiology (Colloca et al., 2017; Gilron et al., 2015). Sensitization is important for neuronal hyperactivity, which is characterized by exaggerated responses to normal stimuli (Campbell and Meyer, 2006; von Hehn et al., 2012). Regulation of various ion channels can contribute to neuronal sensitization in neuropathic pain; however, the ion channel responsible for neuronal hyperactivity remains unknown (Colloca et al., 2017; Gilron et al., 2015). The present study demonstrates that expressional and functional regulation of NALCN can enhance the intrinsic excitability of sensory neurons in neuropathic pain. Specific prevention of up-regulation of NALCN can relieve CCI-induced mechanical allodynia and thermal hyperalgesia *in vivo*, and hyperactivity of sensory neurons is accordingly decreased to normal level. Therefore, NALCN may be a key ion channel for the initiation and maintenance of neuronal sensitization in neuropathic pain and a possible target for the treatment of hyperalgesia.

NALCN is widely expressed in the nervous system. The channel produces a background leak Na^+^ current at the resting membrane potential (RMP) and regulates neuronal excitability and rhythmic behaviors (Ren, 2011). The best-established function of NALCN is maintenance of respiratory rhythms (Lu et al., 2007; Shi et al., 2016). Mice with overall deletion of the NALCN gene develop a severely disrupted respiratory rhythm and die within 24 h after birth (Lu et al., 2007). Mutations of the NCA-1 and NCA-2 gene (homologues of mammalian NALCN) in *Caenorhabditis elegans* lead to abnormalities in locomotion (Jospin et al., 2007). Hypotonia, growth retardation, intellectual disability and apneas induced by central sleep-disordered breathing have been observed in patients with NALCN or UNC-80 mutations (Campbell et al., 2018; Perez et al., 2016). NALCN was also shown to enhance the intrinsic excitability of spinal projection neurons (Ford et al., 2018), which was the first indication that NALCN may contribute to sensory processing. However, the role of NALCN in neuronal sensitization during neuropathic pain is still unclear. The present study indicates that NALCN expression is up-regulated both in the DRG and dorsal spinal cord after CCI, and NALCN current and sensory neurons excitability are increased. Although the percentage of NALCN-positive neurons that also expressed SP increased after CCI in rats, the up-regulation of NALCN was not specific to peptidergic neurons. RNAscope technique was used to confirm that the expression of NALCN mRNA increased commonly in different subtypes of DRG neurons after CCI in mice.

The NALCN complex contains two subunits, UNC79 and UNC80 (Lu and Feng, 2012; Ren, 2011). UNC80 directly interacts with NALCN, and UNC79 indirectly interacts with NALCN *via* UNC80 (Lu and Feng, 2012; Ren, 2011; Wie et al., 2020). The UNC80 and UNC79 subunits are critical for membrane localization and regulation of NALCN (Lu and Feng, 2012; Ren, 2011). However, the specific regulation of NALCN-UNC80-UNC79 complex in neuropathic pain is unclear.

No specific blocker or agonist of NALCN is available. The neuropeptide SP is a modulator of NALCN that can enhance NALCN conductance *via* the neurokinin-1 receptor (NK1R) (Ford et al., 2018; Lu et al., 2009). SP is an 11-amino acid neuropeptide that is synthesized in peptidergic neurons of the DRG and spinal cord (Navratilova and Porreca, 2019). Neuronal terminals in the peripheral and central nervous system can release SP and modulate the transmission of nociceptive signals (Navratilova and Porreca, 2019). Secretion of SP is increased in various pain states including neuropathic pain and inflammatory pain (Milligan and Watkins, 2009; Navratilova and Porreca, 2019; Tiwari et al., 2014). However, the exact target of SP in the enhancement of nociceptive signals remains unclear. SP can elicit pain-related behaviors after intrathecal or local peripheral injection *in vivo (Hylden and Wilcox, 1981; Partridge et al., 1998)*. Here, we showed that the pain-related behaviors induced by intrathecal injection of SP were greatly alleviated after knockdown or knockout of NALCN expression in DRG and dorsal spinal cord, indicating that NALCN may be the molecular target of SP-induced aversive responses.

cAMP (adenosine 3’, 5’ cyclic monophosphate) is a common intracellular second messenger (Skalhegg and Tasken, 2000). The principle intracellular receptor for cAMP in mammals is cAMP-dependent protein kinase (PKA) (Skalhegg and Tasken, 2000). The cAMP/PKA-signaling pathway regulates many cellular processes such as metabolism, cell growth and cell differentiation, as well as ion channel expression and conductivity (Skalhegg and Tasken, 2000). Activation of the cAMP/PKA-signaling pathway contributes to the regulation of ion channels in neuropathic pain (Huang et al., 2012). In this study, the up-regulated expression of NALCN was significantly abolished by intrathecal injection of cAMP or PKA inhibitor at day 3 after CCI, which was accompanied by reduced mechanical allodynia and thermal hyperalgesia *in vivo*. Therefore, the cAMP/PKA-signaling pathway may at least partly contribute to the increased expression of NALCN in neuropathic pain.

The present study used adult rats for *in vivo* behavioral experiments, while neonatal rats were used for patch-clamp recordings. Currently, we can only perform whole-cell recordings on sensory neurons in neonatal spinal cord slices from rats younger than 2 weeks; therefore, rats at postnatal day 5 were subjected to CCI and injected with NALCN-siRNA. Then, whole-cell recordings were performed at day 7-day 8 (2-3 days later). This protocol is also consistent with the behavioral results *in vivo* because the therapeutic effects of NALCN-siRNA are still significant at 2-3 days after injection. Although electrophysiological recordings and therapeutic experiments *in vivo* were not performed in the same rats, the conclusions are reliable because NALCN is widely expressed at both ages (Supplementary fig. 1A), and the neonatal rats exhibited increased NALCN expression in DRG and dorsal spinal cord after CCI that was consistent with the results in adult rats. In addition, the neonatal rats that underwent CCI developed mechanical and thermal hyperalgesia at the age (postnatal day 30) available for behavioral tests. The symptoms of neuropathic pain and neuronal sensitization are thus similar between the different ages.

Regulation of ion channels, including Na_v_ channels, is involved in neuropathic pain (Colloca et al., 2017; Dib-Hajj et al., 2010; Tibbs et al., 2016). We determined that the TTX-R Na_v_ current density is similar between the CCI-ipsilateral and CCI-contralateral sides, indicating that TTX-R Na_v_ may not be involved in the initiation phase of rapid neuronal sensitization in neuropathic pain. However, the contribution of Na_v_ to maintenance of neuronal hyperactivity later cannot be excluded (Colloca et al., 2017; Dib-Hajj et al., 2010). NALCN-siRNA used in this study did not affect TTX-R Na_v_, which can partly confirm its specificity.

Because of the therapeutic role of NALCN for neuropathic pain *in vivo*, a low-dose of NALCN blocker that reverses overactive NALCN may be a novel candidate strategy for the treatment of neuropathic pain. Given the possible toxicity of an NALCN blocker on respiration (Lu et al., 2007), development of an NALCN blocker that is restricted to the peripheral nervous system would be important. We compared the NALCN mRNA level between the spinal cord and heart, and found that NALCN expression in the heart was much lower than that in the nervous system (Supplementary fig. 1B), which is consistent with previous findings (Lee et al., 1999). Low expression level of NALCN in heart may predict a low cardiac toxicity of peripheral restricted NALCN blocker. However, it is still unclear whether NALCN contributes to the cardiac rhythm (Lu and Feng, 2012).

NALCN-siRNA is degraded rapidly after injection (Rao et al., 2009), while the effects of CCI continuously occur. Therefore, CCI-induced mechanical and thermal hyperalgesia are significantly increased on the CCI-ipsilateral side at 5-6 days after injection of NALCN-siRNA. When long-acting AAV-NALCN-shRNA virus or conditional NALCN knockout mice was used, CCI-induced mechanical allodynia and thermal hyperalgesia were completely prevented throughout. Therefore, NALCN may be a key ion channel for the initiation and maintenance of neuronal sensitization in neuropathic pain and a possible target for the treatment of hyperalgesia. Interestingly, increased expression of NALCN can also cause insensitivity to pain (Eigenbrod et al., 2019). Whether high level of NALCN induce hyperalgesia or insensitivity to pain depends on the extent of depolarized RMP. Here we did not elucidate the specific contributions of different neuronal types to NALCN regulation in neuropathic pain.

Although NALCN-siRNA injection decreased the level of NALCN mRNA expression in DRG and dorsal spinal cord, NALCN-siRNA produced no effect on neuronal excitability in electrophysiological recordings and sensory functions *in vivo* without CCI. The relatively low-efficacy of siRNA knockdown may partly explain this result. Additionally, neither AAV-NALCN-shRNA mediated knockdown nor AAV-Cre mediated conditional knockout of NALCN changed the normal sensory functions of rats or mice. Based on above facts, we speculate that conductance of NALCN is small in physiological condition and may limitedly contribute to the normal sensory functions *in vivo*. However, it will be overactive in pathological state, thus contributes to neuronal sensitization in neuropathic pain. Therefore, it is likely that NALCN blocker maybe an underlying therapeutic target for neuropathic pain, while does not affect the normal sensation.

In summary, the present study highlights NALCN as a pivotal ion channel in CCI-induced neuropathic pain and neuronal sensitization. NALCN may be an effective therapeutic target for neuropathic pain.

## Materials and Methods

### Animals

The experimental protocol was approved by the Animal Ethics Committee of West China Hospital of Sichuan University (Chengdu, Sichuan, China) and was conducted in accordance with the Animal Research: Reporting of In Vivo Experiments (ARRIVE) guidelines. Neonatal rats (Postnatal day 5, both sexes), adult male Sprague-Dawley rats (∼8 weeks, 220-250 g), and Nalcn^tm1c(KOMP)Wtsi^ (NALCN^f/f^) mice (Yeh et al., 2017) (∼8 weeks, 20-25 g, both sexes, JAX stock #030718) were maintained under a 12-h (7:00 to 19:00) light/dark cycle at a constant humidity (45-55%) and temperature (22-24°C) with food and water available *ad libitum*. Neonatal rats were kept with the mother. Homozygote wild-type mice (NALCN^+/+^) from the same maternal side (∼8 weeks, 20-25 g, both sexes) were used as control for NALCN^f/f^ mice.

### Injection materials

NALCN-siRNA and control-siRNA with modification of 3’-AlexaFluor488 (QIAGEN, Maryland, USA) were dissolved in RNase-free water (NALCN-siRNA: 5’-GCGAAGAUACCAACGCCAATT-3’; control-siRNA: catalog number 1027419). In vivo SilenceMag™ transfection reagent (OZ Biosciences, Marseille, France) is a highly efficient method dedicated to transfect siRNA into target tissue/cells *in vivo* which combines magnetic nanoparticles and siRNA that will be retained after injection at the magnetically targeted site. In this study, In vivo SilenceMag™ transfection reagent was mixed with NALCN-siRNA or control-siRNA to a concentration of 1 μg/μL 20 min before injection. Then, NALCN-siRNA or control-siRNA was injected into the sciatic nerve (1 μL for neonatal and 2 μL for adult rats) or subarachnoid space (5 μL for neonatal and 10 μL for adult rats). The virus solution of pAAV2-H1-shRNA-(NALCN)-CAG-eGFP (5’-AAGATCGCACAGCCTCTTCAT-3’) or pAAV2-scrambled-CAG-eGFP (5’-GCTCAGTACGATCATACTCAC-3’) (2 x 10^13^ TU/mL) (Taitool Bioscience, Shanghai, China) was unilaterally (left side) injected into the L4-L5 DRG (2 μL per DRG) and subarachnoid space (10 μL) of adult rats. The virus solution of AAV2/9-hSyn-Cre-EGFP-WPRE-pA or AAV2/9-hSyn-EGFP-WPRE-pA (1.2 x 10^13^ V.G./mL) (Taitool Bioscience, Shanghai, China) was unilaterally (left side) injected into the L4-L5 DRG (1 μL per DRG) and subarachnoid space (5 μL) of NALCN^f/f^ and/or wild-type control mice. Substance P (MedChemExpress LLC, Monmouth Junction, NJ, USA) was dissolved in saline and intrathecally injected in volume of 10 μL for adult rats and mice (1 μg/μL). SQ22536 and H-89 dihydrochloride (MedChemExpress LLC, Monmouth Junction, NJ, USA) dissolved in dimethylsulfoxide (DMSO) were intrathecally injected for 3 consecutive days after CCI induction in a volume of 10 μL at concentrations of 1 mM and 10 μM, respectively.

### CCI model

A CCI-induced neuropathic pain model was established (Bennett and Xie, 1988). Briefly, the rats/mice were anesthetized by 2-3 % sevoflurane. Then, the left sciatic nerve at the level of the mid-thigh was exposed and loosely ligated with 4-0 chromic gut sutures at four sites for adult rats or 5-0 chromic gut sutures at three sites for neonatal rats or adult mice. The right sciatic nerve was exposed but not ligated and is referred to as the CCI-contralateral side.

### Sciatic nerve, DRG and intrathecal injection in rats

Injection of the sciatic nerve was performed at the proximal site of ligation with a glass micropipette connected to a Hamilton syringe. DRG injection was performed as described previously (Yu et al., 2016). Briefly, adult rats were anesthetized with 2-3% sevoflurane. DRG (left L4-L5) were surgically exposed and the pAAV2-H1-shRNA-(NALCN)-CAG-eGFP or pAAV2-scrambled-CAG-eGFP virus (2 x 10^13^ TU/ml) was unilaterally injected into the DRG (2 μL per DRG) at a rate of 0.2 μL/min with a glass micropipette connected to a Hamilton syringe controlled by stereotaxic syringe holder (SYS-Micro4, WPI, Sarasota, FL, USA). The needle was removed 10 min later. Rats were used for experiments 4 weeks later. Intrathecal injection was performed in anesthetized rats (Mestre et al., 1994) except for SP injection to exclude the effects of anesthesia on SP-induced pain behaviors. The rats were restrained in horizontal position, and a 25-G needle connected to a Hamilton syringe was inserted perpendicular to the vertebral column between the L4 and L5 lumbar vertebrae. Sudden movement of tail was used as an indicator of successful puncture into the subarachnoid space. The syringe needle was held in place for at least 30 s to avoid any outflow of the solution. A magnet (SilenceMag™ kit) was placed at the body surface corresponds to the targeted site for at least 1 hour after injection of NALCN-siRNA and control-siRNA.

### Cre-dependent conditional knockout of NALCN in DRG and spinal cord neurons in mice

In brief, C57BL6 mice (∼8 weeks, 20-25 g) containing loxP sites inserted to flank NALCN protein coding regions (NALCN^f/f^) or wild-type littermates (NALCN^+/+^) were anesthetized with 2-3% sevoflurane. The mice littermates then received both DRG (left L4-L5) and intrathecal injection of AAV2/9-hSyn-Cre-EGFP-WPRE-pA or AAV2/9-hSyn-EGFP-WPRE-pA. Cre-mediated excision of the floxed region produces no detectable transcript which can be determined by RT-PCR. Mice were used for experiments 4 weeks later.

### Behavioral tests

Von Frey filaments and the Hargreaves test (heat) were used to measure mechanical allodynia and thermal hyperalgesia (Hargreaves et al., 1988; Singh et al., 2009). For the Von Frey test, rats/mice were placed in a Plexiglas chamber on an elevated metal mesh floor and habituated for 30 min. The plantar surface of the hind limb was stimulated with a series of Von Frey filaments (0.008, 0.02, 0.04, 0.16, 0.4, 0.6, 1, 1.4, 2, 4, 6, 8, 10 and 15 g). The absolute withdrawal threshold was determined using the up-and-down method (Singh et al., 2009). For the Hargreaves test, the rats/mice were placed in the same chamber, which was placed on a transparent glass plate. The plantar surface of the hind limb was exposed to a beam of radiant heat (∼55°C for rats and 35°C for mice) through the transparent glass. Thermal withdrawal latencies were recorded, and the stimulus cut-off time was set at 15 s to prevent injury. Measurements were repeated three times with a 10-min interval between measurements, and the mean latency was calculated. All behavioral tests were performed from 9:00 to 12:00 from the post-CCI day 1 to 14. After injection of SQ22536 and H89 dihydrochloride, behavioral tests were performed 4 h later. For SP-induced pain behaviors, the cumulative time of biting and scratching episodes directed toward the lumbar and caudal region of spinal cord were measured with a stopwatch during the first period of 5 min after intrathecal injection of SP. The individual experimenter was blinded to the treatment group of the rats.

### Real-time PCR

Total RNA was isolated using the Eastep^®^ Super RNA extraction kit (Promega, Shanghai, China). Reverse transcription was performed with a GoScript™ Reverse Transcription Kit (Promega, Shanghai, China). RT-PCR was performed using GoTaq® qPCR Master Mix (Promega, Shanghai, China) and specific primers (Sangon Biotech, Shanghai, China) according to the manufacturer’s protocol. Gene expression was normalized to the GAPDH level. The primers used to detect NALCN and GAPDH mRNA were as follows:

NALCN forward (5’-GTCCTGACGAATCTCTGTCAGA-3’), NALCN reverse (5’-CTGAGATGACGCTGATGATGG-3’); GAPDH forward (5’-GACATGCCGCCTGGAGAAAC-3’), GAPDH reverse (5’-AGCCCAGGATGCCCTTTAGT-3’).

### Immunofluorescence staining

Rats were anesthetized with ketamine/xylazine (60/10 mg/kg) and transcardially perfused with ice-cold Ringer’s solution followed by 4% paraformaldehyde. The spinal cord and DRG were removed and stored in a 4% paraformaldehyde solution overnight, followed by incubation in 30% sucrose for 1 day (for the DRG) or 2 days (for the spinal cord). Transverse sections (12 μm) were cut using a freezing microtome (CM1850; Leica, Buffalo Grove, IL, USA). Sections were incubated at 4°C overnight with the following primary antibodies: NALCN (1:400, rabbit, Alomone Labs, Cat#:ASC022), TRPV1 (1:800, mouse, Abcam, Cat#:ab203103), SP (1:800, mouse, Millipore, Cat#:MAB356), neurofilament (1:800, mouse, Millipore, Cat#:MAB5262), and IB4-FITC-conjugated 488 (1:400, Life, Cat#:I21411). The sections, except for slices incubated with IB4-FITC-conjugated 488 antibodies, were incubated with secondary antibodies for 2 h: Alexa Fluor 488 goat anti-mouse IgG (1:400, Millipore), or Fluor 594 goat anti-rabbit IgG (1:400, Millipore), except for IB4. Specificity of the NALCN primary antibody was validated by pre-incubating with the antigen (available from the manufacturer) as presented in Supplementary Fig. 5B. Fluorescent images were acquired using Zeiss AxioImager Z.2 (Guangzhou, Guangdong, China).

### Preparation of spinal cord slices

Rats (P7-P8) were anesthetized with ketamine/xylazine (60/10 mg/kg). Then, the lumbar enlargement of the spinal cord was obtained, fixed in agarose gel (1.8%) and mounted on a vibratome (VT1000 A; Leica). Sagittal spinal cord slices (270 μm in thickness) were cut and incubated in external solution containing (in mM): 130 NaCl, 3 KCl, 2 MgCl_2_, 2 CaCl_2_, 1.25 NaH_2_PO_4_, 26 NaHCO_3_, and 10 glucose at 35°C for 45 min and then held at room temperature (24-26°C). The incubation solution was aerated with 95% O_2_/5% CO_2_. After incubation, spinal cord slices were mounted in the recording chamber for recordings at room temperature.

### Acute isolation of DRG neurons

Acute isolated DRG neurons were prepared as follows (Zheng et al., 2007). Briefly, rats at age of P7-P8 were anesthetized with ketamine/xylazine (60/10 mg/kg). DRGs were removed and digested sequentially in 1% papain for 25 min and in 1% collagenase I for 20 min at 37°C. Isolated neurons were transferred to glass coverslips coated with poly-D-lysine and laminin. The neurons were incubated with DMEM aerated with 95% O_2_/5% CO_2_. Patch clamp recordings were performed after the neurons attached to the coverslips (∼2 h).

### Patch-clamp recordings

Spinal cord slice or acute isolated DRG neurons were mounted in a recording chamber and submerged in continuously perfused incubation solution (∼2 mL/min), bubbled with 95% O_2_/5% CO_2_. The incubation solution contained (in mM): 140 NaCl, 3 KCl, 2 MgCl_2_, 2 CaCl_2_, 10 HEPES, and 10 D-glucose. DRG and dorsal spinal cord (lamina I-II) neurons were visualized and identified by shape using infrared differential interference contrast microscopy. Electrophysiological recordings were conducted using an Axopatch 700B amplifier and a Digidata 1440 digitizer linked to a computer running pClamp 10.2 software (Molecular Devices, Sunnyvale, CA, USA). Recordings were sampled at 20 kHz and filtered at 10 kHz. All recordings were performed in the whole-cell configuration. The resistance of patch electrodes was 3-6 MΩ. Current-clamp recordings were performed to record APs using an internal solution containing (in mM): 120 KCH_3_SO_3_, 4 NaCl, 1 MgCl_2_, 0.5 CaCl_2_, 10 HEPES, 10 EGTA, 3 Mg-ATP, and 0.3 GTP-Tris, pH 7.35. APs were recorded with perfusion of 10 μM bicuculline and 100 μM picrotoxin to block possible effects from synaptic and/or extrasynaptic GABAergic inputs. For the recordings performed in current-clamp mode, the rheobase was determined by applying short depolarizing current steps (5 pA steps with a duration of 100 ms) until a single AP was generated. The firing frequency was examined using sustained depolarizing current steps (10 pA steps with a duration of 1000 ms). Voltage-clamp recordings were established to record NMDG- (140 mM), Gd^3+^- (50 μM), and SP-sensitive (10 μM) holding currents using an internal solution containing (in mM): 110 CsCH_3_SO_3_, 9 NaCl, 1.8 MgCl_2_, 4 Mg-ATP, 0.3 Na-GTP, 0.09 EGTA, 0.018 CaCl_2_, 9 HEPES, and 10 TEA-Cl, pH 7.38. The series resistance was compensated by ∼70%-75%, and data were rejected when the series resistance exceeded 20 MΩ. To block Na_v_ and background potassium currents, 25 mM TEA, 5 mM 4-AP, 500 nM TTX and 100 μM CsCl were added to the incubation solution.

### RNAscope in situ hybridization

RNAscope *in situ* hybridization was performed as described previously (Liao et al.). The sequences of the target probes, pre-amplifier, amplifier, and label probes were commercially available (Advanced Cell Diagnostics, CA, US). The probes included *Nalcn* (Cat#415161), *Nefh* (Cat#443671), *Tac1* (Cat#410351), *Th* (Cat#317621), *P2rx3* (Cat#521611) and *Rbfox3* (Cat#313311). Fluorescence images were acquired with a NIKON A1R^+^ two-photon confocal scanning microscope (Shanghai, China) and were processed with ImageJ software (NIH, Bethesda, MD, US). Quantifications of expression of NALCN mRNA in all DRG neurons (*Rbfox3*) or different subtypes of DRG neurons (*Nefh*, *Tac1*, *Th* and *P2rx3*) was present by mean fluorescence intensity of area of total DRG neurons or every positive neuron, respectively.

### Statistical analysis

Data were expressed as means ± SEM and statistical analyses were performed using GraphPad Prism 8.0 software (GraphPad Software, CA, USA). Paired/unpaired Student’s *t*-tests or Mann-Whitney U test were used for comparisons of parametric distribution data or nonparametric distribution data between two groups, respectively. The exact analysis used for each comparison was described in figure legends. Time course data of behavioral tests were analyzed using a two-way analysis of variance (ANOVA) with repeated measures followed by a Bonferroni *post-hoc* test. *P* < 0.05 was considered statistically significant.

## Acknowledgments

None.

## Funding

This study was supported by the grant No. 81974164 and 81771486 (to C.Z.), No. 81600918 (to P.L.) and 81571300 (to J.L.) from National Natural Science Foundation of China (Beijing, China).

## Author contributions

C.Z. and H.H. conceived the study. C.Z., D.Z. and W.Z. designed the study. D.Z., W.Z., J.L., M.O., D.L., S.B., J.L., Y.C., P.L., J.S. and X.C. performed the experiments and analyzed the data. C.Z., H.H., D.Z. and W.Z. wrote the manuscript.

## Conflict of interest

The authors declare that they have no competing interests.

## Data availability

This study includes no data deposited.

## Supplementary Materials

**Supplementary Fig. 1.**
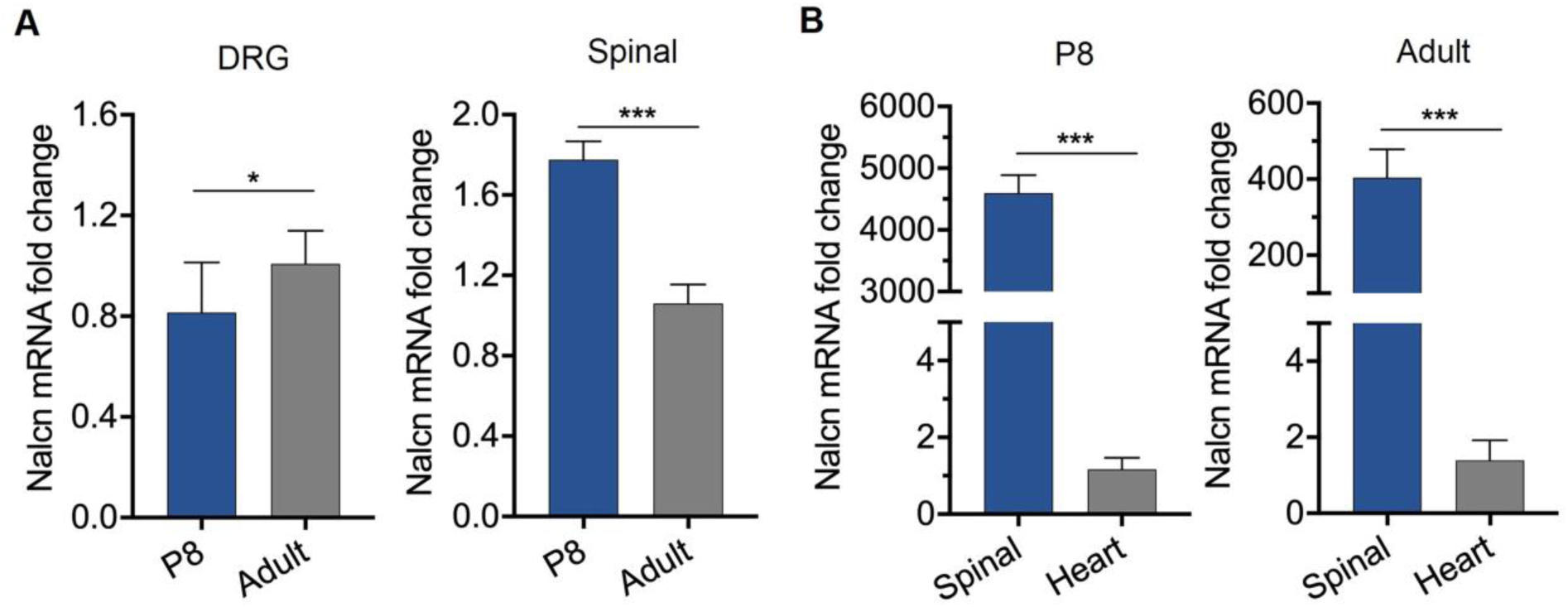
NALCN mRNA level is age-dependent and tissue-dependent. (**A**) NALCN mRNA level was higher in DRG (left panel, n = 9 for P8, n = 12 for Adult) but lower in spinal cord (right panel, n = 9 for P8, n = 12 for Adult) of adult rats compared to the rats at age of P8. (**B**) NALCN mRNA level was much lower in heart compared to dorsal spinal cord of rats at both ages of P8 (left panel, n = 6) or adult (right panel, n = 6). Data are present as mean ± SEM. A by unpaired two-tailed student’s t-test; B by paired two-tailed student’s t-test. \**P* < 0.05 and \*\*\**P* < 0.001.

**Supplementary Fig. 2.**
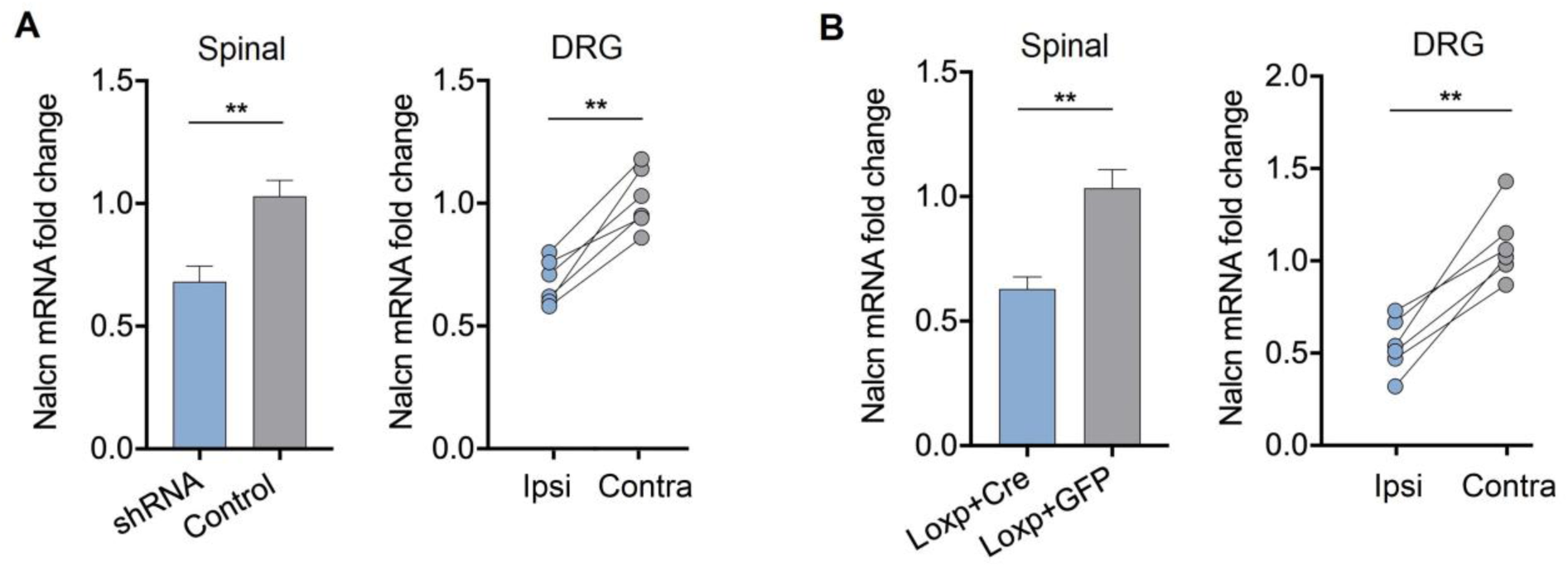
The knockdown/knockout efficiency of NALCN was confirmed by RT-PCR. (**A**) The level of NALCN mRNA was significantly decreased in dorsal spinal cord (left panel, n = 6) of rats 4 weeks after intrathecal injection of AAV-NALCN-shRNA as compared to AAV-scrambled-shRNA. The level of NALCN mRNA was significantly decreased in DRG 4 weeks after DRG injection of AAV-NALCN-shRNA (right panel, n = 6) but not AAV-scrambled-shRNA. (**B**) The level of NALCN mRNA was significantly decreased in dorsal spinal cord (left panel, n = 6) of NALCN^f/f^ mice 4 weeks after intrathecal injection of AAV-Cre as compared to AAV-GFP. NALCN mRNA was significantly decreased in DRG of NALCN^f/f^ mice 4 weeks after DRG injection of AAV-Cre (right panel, n = 6) but not AAV-GFP. Data are present as mean ± SEM. A left panel and B left panel by unpaired two-tailed student’s t-test; A right panel and B right panel by paired two-tailed student’s t-test. ** *P* < 0.01.

**Supplementary Fig. 3.**
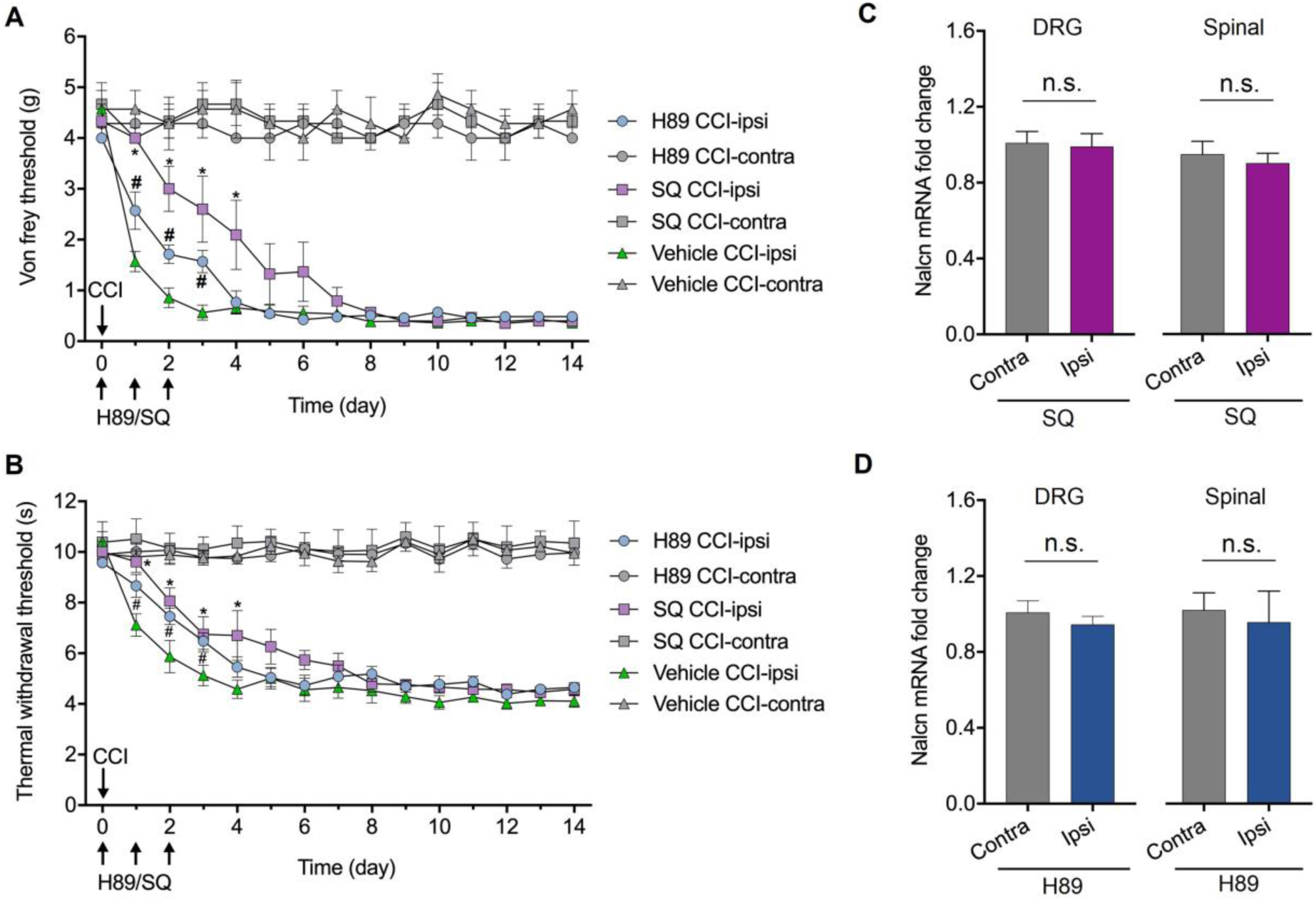
cAMP-PKA pathway regulates the expression of NALCN after CCI. (**A** and **B**) Intrathecal injection of SQ22536 (cAMP inhibitor) or H89 (PKA inhibitor) alleviated the CCI-induced mechanical allodynia (A, n = 6 for SQ22536 group, n=7 for H89 group, ^#^*P* < 0.05 vs. CCI-ipsi + vehicle, \**P* < 0.05 vs. CCI-ipsi + vehicle) and thermal hyperalgesia (B, n = 6 for SQ22536 group, n = 7 for H89 group, ^#^ *P* < 0.05 vs. CCI-ipsi + vehicle, \**P* < 0.05 vs. CCI-ipsi + vehicle). (**C**) NALCN mRNA was not increased at day 3 after CCI by SQ22536 in DRG (left panel, n = 6) and dorsal spinal cord (right panel, n = 6) in the CCI-ipsilateral side compared to the CCI-contralateral side. (**D**) NALCN mRNA was not increased at day 3 after CCI by H89 in DRG (left panel, n=7) and dorsal spinal cord (right panel, n=7) in the CCI-ipsilateral side compared to the CCI-contralateral side. Data are present as mean ± SEM. A and B by two-way ANOVA; C and D by paired two-tailed student’s t-test. n.s.: no significance.

**Supplementary Fig. 4.**
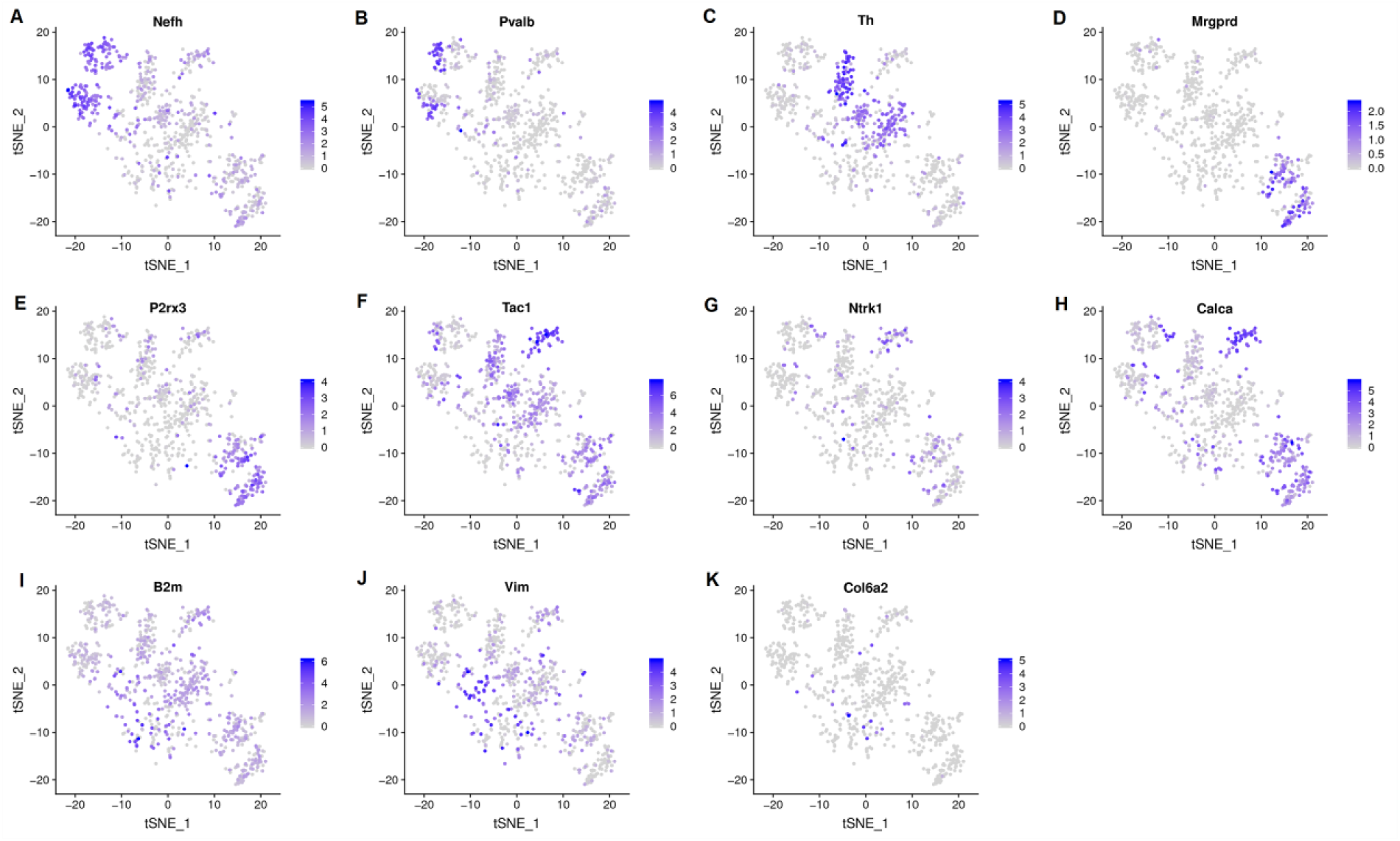
The expression profile of well-known markers of the five clusters was showed. t-Distributed stochastic neighbor embedding identified 11 major cell populations showing the expression of representative well-known cell-type-specific marker genes. Numbers reflect the number of UMI detected for the specified gene for each cell.

**Supplementary Fig. 5.**
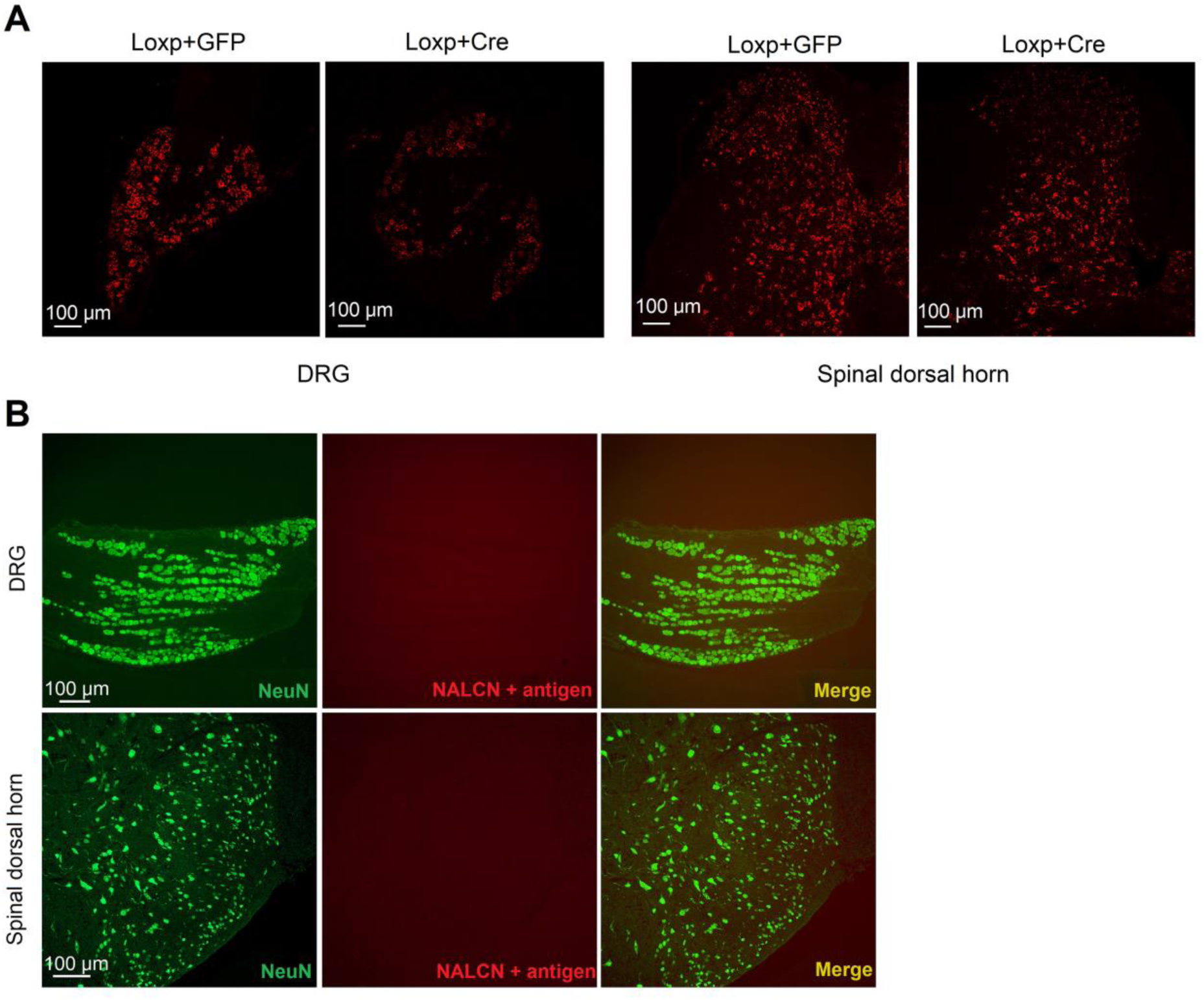
Representative images of the validation of Cre-dependent conditional knockout of NALCN and the specificity of NALCN antibody in DRG and spinal cord neurons. (**A**) Representative images showed the expression of NALCN mRNA in DRG and spinal cord neurons from NALCN^f/f^ mice that received AAV-Cre or AAV-GFP using RNAscope. NALCN positive fluorescence was decreased in DRG (left panel) and spinal cord (right panel) in NALCN^f/f^ mice that received AAV-Cre as compared to AAV-GFP. (**B**) Representative images showed that the specificity of NALCN antibody was validated in DRG and spinal cord of rats by immunofluorescence staining, related to Figure 1. NALCN positive fluorescence was almost diminished in DRG (top panel) and spinal cord (bottom panel) when the antibody pre-incubated with NALCN antigen.

**Supplementary Fig. 6.**
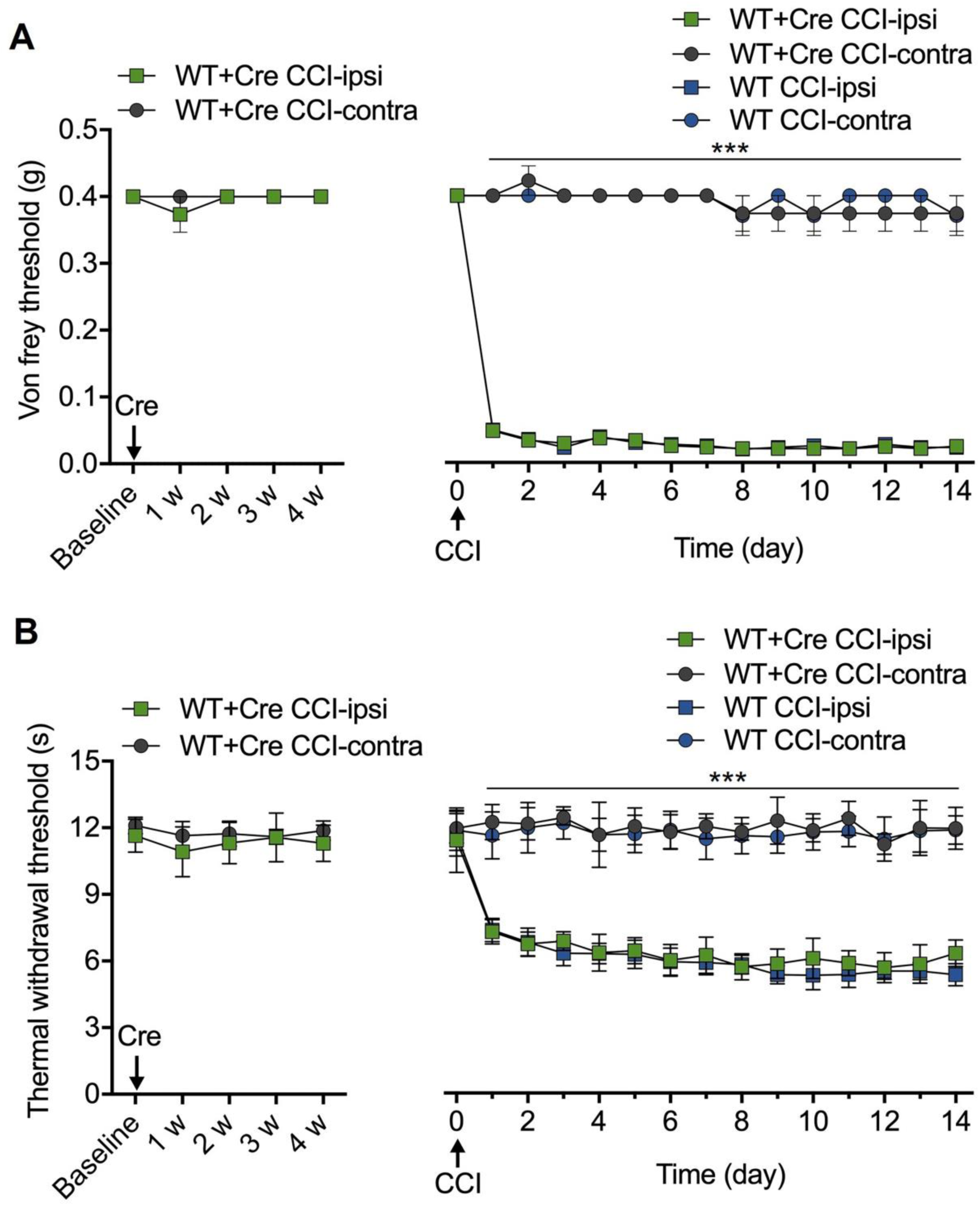
Injection of AAV2/9-hSyn-Cre-EGFP-WPRE-pA produce no effects on the CCI-induced mechanical allodynia and thermal hyperalgesia in wild-type (NALCN^+/+^) mice. (**A** and **B**) After DRG and intrathecal injection of AAV2/9-hSyn-Cre-EGFP-WPRE-pA, normal sensation including mechanical allodynia (A, left panel, n = 9 for WT + Cre group) and thermal hyperalgesia (B, left panel, n = 9 for WT + Cre group) was unchanged when tested once a week after injection. Mechanical allodynia (A, right panel, n = 9 for WT + Cre group, n = 8 for WT group) and thermal hyperalgesia (B, right panel, n = 9 for WT + Cre group, n=8 for WT group) were developed in CCI-ipsilateral side since the first day after CCI until at least day 14 in WT mice or WT mice that received DRG and intrathecal injection of AAV-Cre. Data are present as mean ± SEM. A and B by two-way ANOVA; *** *P* < 0.001. n.s.: no significance.

